# Dynamics of transcription elongation are finely tuned by dozens of regulatory factors

**DOI:** 10.1101/2021.08.15.456358

**Authors:** Mary Couvillion, Kevin M. Harlen, Kate C. Lachance, Kristine L. Trotta, Erin Smith, Christian Brion, Brendan M. Smalec, L. Stirling Churchman

## Abstract

Understanding the complex network and dynamics that regulate transcription elongation requires the quantitative analysis of RNA polymerase II (Pol II) activity in a wide variety of regulatory environments. We performed native elongating transcript sequencing (NET-seq) in 41 strains of *S. cerevisiae* lacking known elongation regulators, including RNA processing factors, transcription elongation factors, chromatin modifiers, and remodelers. We found that the opposing effects of these factors balance transcription elongation dynamics. Different sets of factors tightly regulate Pol II progression across gene bodies so that Pol II density peaks at key points of RNA processing. These regulators control where Pol II pauses with each obscuring large numbers of potential pause sites that are primarily determined by DNA sequence and shape. Genes that are sensitive to disruptions in transcription elongation tend to couple changes in Pol II pausing and antisense transcription to transcription output. Our findings collectively show that the regulation of transcription elongation by a diverse array of factors affects gene expression levels and co-transcriptional processing by precisely balancing Pol II activity.

## INTRODUCTION

Transcription is a highly regulated and conserved process that consists of three phases: initiation, elongation, and termination (Shandilya and Roberts, 2012; Svejstrup, 2004). Post-initiation regulation is critical for co-transcriptional RNA processing, shaping the chromatin landscape, and preventing run-on transcription into downstream genes (Herzel et al., 2017; Holmes et al., 2015; Proudfoot et al., 2002; Rando and Winston, 2012). Transcription elongation is controlled across gene bodies by a wide variety of factors, including transcription factors, chromatin modifiers, chromatin assembly factors and chaperones, RNA processing factors, and histone variants. Understanding how these factors act separately and in concert to influence RNA polymerase II (Pol II) activity will shed light on how transcription elongation and co-transcriptional processes are coordinated.

Transcription is a discontinuous process: periods of productive elongation are frequently interrupted by pauses. Pol II pausing was first observed *in vitro* in *E. coli* polymerase transcribing the *lac* operon and lambda DNA (Dahlberg and Blattner, 1973; Gilbert et al., 1974; Kassavetis and Chamberlin, 1981; Kingston and Chamberlin, 1981; Lee et al., 1976; Maizels, 1973). Observations of Pol II pausing *in vivo* provided the first evidence of promoter-proximal pausing (Gariglio et al., 1981). These findings were extended by chromatin immunoprecipitation (ChIP) studies, which identified paused polymerase near the 5’ ends of certain *Drosophila* and mammalian genes (Bentley and Groudine, 1986; Eick and Bornkamm, 1986; Gilmour and Lis, 1986; Krumm et al., 1992; Nepveu and Marcu, 1986; Rougvie and Lis, 1988; Spencer and Groudine, 1990; Strobl and Eick, 1992). Promoter-proximal pausing is now known to occur genome-wide and has been visualized by high-resolution ChIP-seq (ChIP-exo) experiments, which have pinpointed Pol II accumulation 50 bp downstream of transcription start sites (TSSs) in the majority of human genes (Venters and Pugh, 2013). The advent of high-throughput and high-resolution sequencing technologies have led to the development of sequencing methods such as native elongating transcript sequencing (NET-seq) and precision run-on sequencing (PRO-seq) that measure Pol II density genome-wide at nucleotide resolution. Collectively, these techniques have highlighted the control of transcription elongation by regulatory factors. These approaches and other nascent RNA sequencing methods visualize the production of transcripts from RNA polymerases across the genome (Churchman and Weissman, 2011; Core et al., 2008; Kwak et al., 2013; Mayer et al., 2015; Nojima et al., 2015; Schwalb et al., 2016), and therefore are capable of revealing the immediate and direct effects of a perturbation on transcription. In addition, these assays capture unstable transcripts such as enhancer RNAs and antisense RNAs, which are critical to transcription regulation but invisible to other techniques (Faghihi and Wahlestedt, 2009; Hou and Kraus, 2020). The strand-specificity and high resolution of these methods are transforming our understanding of transcription dynamics and regulation.

NET-seq, PRO-seq, and other high-resolution methods have revealed regions of high Pol II density, such as promoter proximal pausing, and specific sites of Pol II pausing within the region and throughout the gene body (Churchman and Weissman, 2011; Ferrari et al., 2013; Kindgren et al., 2020; Kwak et al., 2013; Larson et al., 2014; Mayer et al., 2015; Nojima et al., 2015; Vvedenskaya et al., 2014; Weber et al., 2014). These Pol II pause sites are reminiscent of pausing observed at single nucleotide positions *in vitro* (Churchman and Weissman, 2011; Kassavetis and Chamberlin, 1981; Kwak et al., 2013; Schwalb et al., 2016). In yeast, Pol II pause sites occur frequently (Churchman and Weissman, 2011), arising from intrinsic properties of the polymerase itself, interactions with the DNA template and specific sequence motifs, and the presence of bound proteins (e.g. histones and transcription factors) (Gajos et al., 2021; Herbert et al., 2006; Hodges et al., 2009; Kassavetis and Chamberlin, 1981; Kireeva and Kashlev, 2009; Kireeva et al., 2005; Mayer et al., 2015; Noe Gonzalez et al., 2021; Shaevitz et al., 2003). In yeast, Pol II tends to pause at an adenine and just before the nucleosome dyad (Churchman and Weissman, 2011; Hodges et al., 2009), the point of the strongest DNA–histone contacts (Hall et al., 2009). The presence of elongation factors changes the duration and position of Pol II pauses *in vitro* and *in vivo*. For example, in the absence of Dst1, the yeast homolog of the elongation factor TFIIS, ∼75% of Pol II pauses are shifted 5–18 bp downstream (Churchman and Weissman, 2011). *In vitro,* the TFIIS bacterial homolog, NusG, changes the duration of RNA polymerase pausing (Herbert et al., 2010).

Regions or peaks of high Pol II density, such as promoter proximal pauses, are created in part by a high density of pause sites that together create barriers to elongation and provide an opportunity for regulation and coordination of co-transcriptional events (Bentley, 2014; Mayer et al., 2017; Noe Gonzalez et al., 2021; Rougvie and Lis, 1988). For example, prominent peaks of Pol II density occur at the boundaries of retained exons and near polyadenylation [poly(A)] sites (Harlen et al., 2016; Kwak et al., 2013; Mayer et al., 2015; Nojima et al., 2015). Myriad factors control Pol II peaks *in vivo*. Loss of Rtt103, a termination factor, causes a dramatic peak in Pol II density directly downstream of poly(A) sites (Harlen et al., 2016).

The full range of Pol II pausing preferences and behavior is relatively unknown, and the degree to which pausing can be suppressed or enhanced by perturbing the transcription regulatory network has yet to be determined. For example, we do not know which elongation factors are responsible for Pol II pausing, nor which features of chromatin determine pausing location. Moreover, the connection between Pol II pausing and overall gene expression levels remains poorly understood. To answer these questions, we need to study pausing in diverse regulatory landscapes and functionally characterize factors involved in the regulation of transcription elongation.

Pol II transcribes much of the genome in all eukaryotes, yet only a fraction of its transcripts mature into stable, protein-coding RNA products (Bertone et al., 2004; Cheng et al., 2005; David et al., 2006; Hangauer et al., 2013; Kapranov et al., 2007; Mercer et al., 2011; Nagalakshmi et al., 2008; Steinmetz et al., 2006). A major contributor to unstable noncoding RNA products is antisense transcripts, i.e., RNAs transcribed from the strand opposite the sense strand of a protein-coding gene. Originally identified in bacteria (Spiegelman et al., 1972), antisense transcripts were soon discovered in eukaryotes as well (Anderson et al., 1981; Bibb et al., 1981). Since its discovery, antisense transcription has been detected opposite the vast majority of annotated genes in yeast (Xu et al., 2011).

Although a general, genome-wide function has not been identified, antisense transcription has been proposed to regulate transcription factor recruitment and transcriptional repression (Donaldson and Saville, 2012; Gullerova and Proudfoot, 2010; Lenstra et al., 2015; Scruggs et al., 2015). Importantly, misregulation of antisense transcription can alter the transcriptional landscape and chromatin architecture of cells (Gullerova and Proudfoot, 2010; Marquardt et al., 2014; Pelechano et al., 2013; Wei et al., 2011). To better understand pervasive antisense transcription and its role in regulation, we need to determine whether it is tunable by regulatory factors. If antisense transcription levels are tightly controlled, it is possible that antisense transcription could be manipulated in order to contribute to overall transcription regulation.

To gain insight into the regulation of the production of coding and non-coding transcripts by Pol II, we used NET-seq to analyze 41 *S. cerevisiae* mutant strains lacking known elongation regulators. We investigated how each factor regulates nascent transcription, production of antisense transcripts and pausing across gene bodies. Metrics describing each transcription phenotype span a broad dynamic range with wild-type activity lying near the center. The loss of each factor revealed distinct sets of pause sites that we used to create machine learning models of Pol II pausing, highlighting which genomic features can predict pause positions. Finally, we investigated genes that are frequently regulated across the strains and identified relationships between transcription elongation dynamics, antisense transcription and transcriptional output. Together, our results show that Pol II dynamics are determined by the contrasting impacts of regulatory factors that, in turn, impact the overall production of RNA transcripts.

## RESULTS

### Reverse genetic screen for transcription regulators

To obtain insight into the transcription regulatory network of *S. cerevisiae*, we individually deleted 41 known transcription elongation regulators, including RNA processing factors, transcription elongation factors, histone variants, chromatin modifiers, and chromatin remodelers and chaperones, and assessed the transcriptional effects of each deletion using NET-seq (**Figure 1A**). The wild-type transcription baseline was established using four biological replicates of wild-type cells; the results from the replicates were highly correlated (R^2^ ≥ 0.97; **Figure S1A**). All mutant strains were analyzed in at least biological duplicate. Results from strain replicates were highly correlated (R^2^ ≥ 0.75; **Table S1**). Importantly, all replicates were performed at different times, by different researchers, and in different strain isolates, demonstrating the reproducibility of our results.

**Figure 1.**
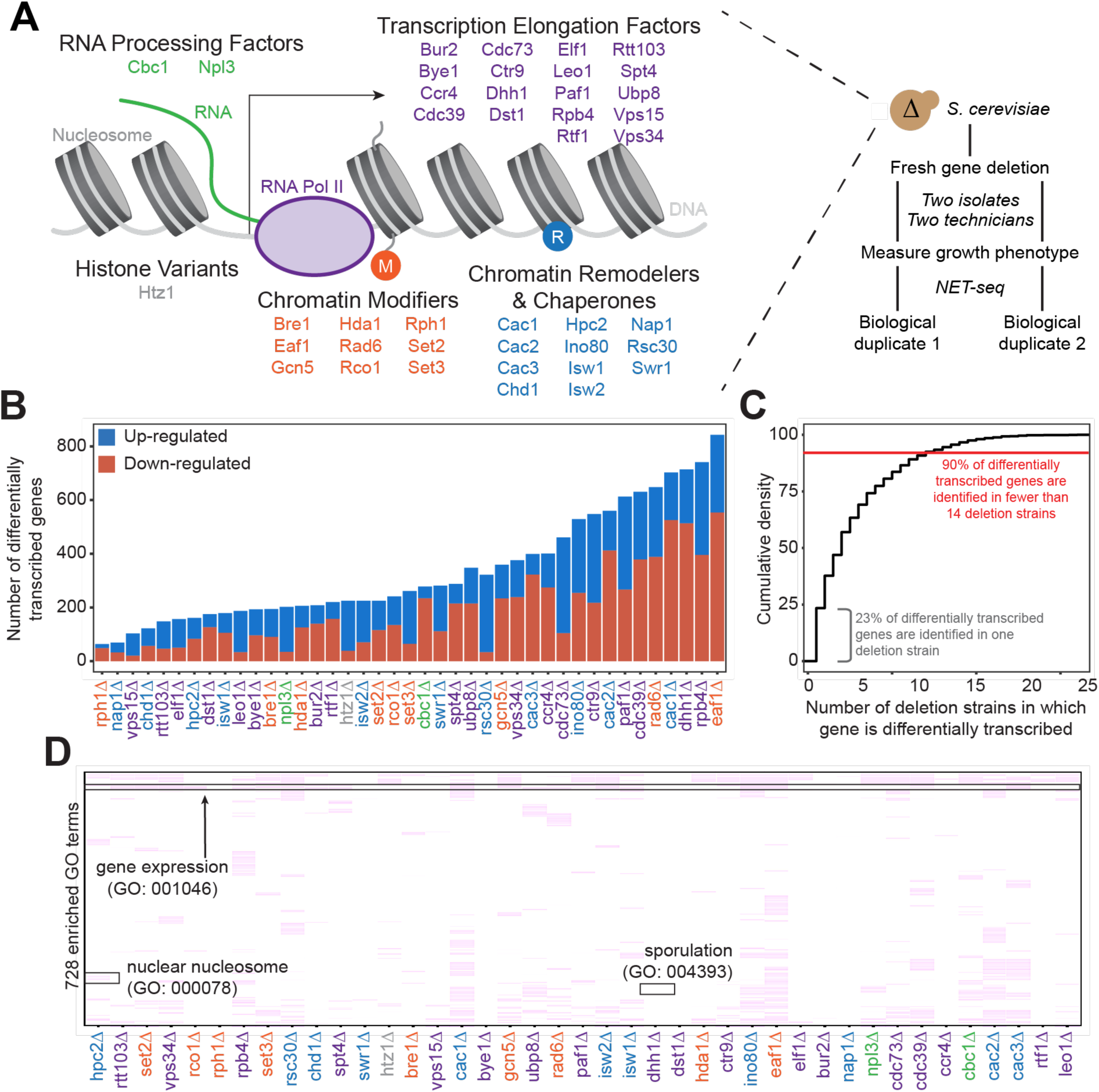
Gene expression is affected differently when transcription regulatory proteins are knocked out, both at the level of individual genes and gene ontology. (A) As Pol II transcribes along a chromatinized template, a complex network regulates eukaryotic transcription elongation. Factors analyzed in the reverse genetic screen are listed and grouped by function: RNA processing factors (*green*), transcription elongation factors (*purple*), histone variants (*grey*), chromatin modifiers (*orange*), and chromatin remodelers and chaperones (*blue*). Colors of factors consistent throughout figures. Each of these factors were deleted to conduct a reverse genetic screen in *S. cerevisiae*. For each deletion strain, a fresh gene deletion was conducted in two isolates by two technicians. After a growth phenotype was measured, NET-seq was performed in at least two biological replicates. (B) Number of differentially up- (*blue*) and down-regulated (*red*) genes vary widely across deletion strains. (C) Cumulative density plot illustrating that 27% of DE genes are only differentially transcribed in one strain, with 90% of DE genes differentially transcribed in 10 strains or fewer. There are several genes that are expressed in 24 out of 31 deletion strains for which there were sufficient replicates to conduct differential expression analysis. (D) 303 gene ontology terms are enriched (*purple*) in at least one of the strain’s differentially transcribed genes; if a GO term is not enriched in a deletion strain, the heatmap tile is white. Both axes are hierarchically clustered to group those deletion strains that share enriched ontologies. Differential transcription and GO term enrichment analyses conducted only on deletion strains with biological replicates.

### Nascent gene expression is uniquely disrupted across deletion strains

Because all of the factors examined in our screen play roles in transcription regulation, we first sought to determine whether each factor regulates different sets of genes, or whether modifications of the transcriptional regulation network affect the transcription of overlapping sets of genes. Based on NET-seq data, we assessed the role of each factor in regulating nascent transcription, a more direct measurement of transcriptional phenotype than can be obtained from RNA-seq data. Interestingly, in some strains (e.g., *rph1Δ* and *nap1Δ*), very few genes were transcribed at significantly altered levels relative to the wild-type, whereas in others (e.g. *eaf1Δ* and *dhh1Δ*), over 15% of all protein-coding genes were differentially transcribed (**Figures 1B & S1B; Table S2**).

We then investigated the degree to which differentially transcribed genes were shared across mutant strains. Over 90% of differentially transcribed genes were identified in fewer than 11 deletion strains, and 27% were differentially transcribed in only a single strain (**Figure 1C**). Only a few genes had altered expression in the majority of deletion strains; some of these, such as *HSP26* are involved in stress responses, and their regulation may represent the cell’s reaction to losing key transcription regulators.

We postulated that genes that were differentially transcribed in many strains might have unique features that sensitize them to disruptions in the transcription elongation regulation network. We identified 155 genes up-regulated and 173 genes down-regulated in at least 8 strains. Gene Ontology (GO) enrichment analysis did not reveal significant enrichment of common functions, although genes involved in metabolism, ion transport, and cell wall/membrane components and assembly were weakly enriched (< 6.5-fold) (Anders and Huber, 2010; Ashburner et al., 2000; Mi et al., 2019; The Gene Ontology Consortium, 2019). Notably from the standpoint of transcription regulation, these genes had a greater propensity to have TATA-containing promoters (p < 0.001 by Chi-squared test) and tended to be shorter than the average gene (p < 0.001 by Student’s t-test, **Figure S1C-D**). This is consistent with the fact that 20% of all yeast genes whose promoters contain TATA boxes are associated with stress responses and are under high degrees of regulatory control (Basehoar et al., 2004). These results suggest that gene length and promoter composition may predispose genes to transcriptional changes upon perturbations to the transcription regulatory network

We next asked whether certain biological functions or pathways were commonly affected across the deletion strains using GO enrichment analysis (**Figure 1D; Table S3**) (Anders and Huber, 2010; Ashburner et al., 2000; Mi et al., 2019; The Gene Ontology Consortium, 2019). Over 90% of GO pathways enriched among the differentially transcribed genes were identified in fewer than nine deletion strains, with 40% identified in a single strain, emphasizing the largely distinct responses to loss of each factor (**Figure S1E**). GO enrichments were not particularly strong or specific overall (**Table S3**); however, we did detect enrichment of some pathways consistent with the known functions of certain factors. Upon deletion of *HPC2*, which encodes a subunit of the HIR nucleosome assembly complex involved in the regulation of histone gene transcription (Formosa et al., 2002; Prochasson et al., 2005; Xu et al., 1992), the term ‘nuclear nucleosome’ (GO:000078) was significantly enriched (36.5 fold enrichment, FDR = 0.023) among differentially transcribed genes. Upon deletion of *DHH1*, differentially transcribed genes were enriched for functions related to the sporulation pathway (GO:004393, 3.0 fold enrichment, FDR < 0.001), which is nonfunctional in *DHH1*-null mutants (Enyenihi and Saunders, 2003; Moriya and Isono, 1999). Interestingly, in half of all deletion strains, differentially transcribed genes were enriched for functions related to regulation of gene expression (GO:001046), suggesting feedback loops between perturbation of genes that regulate gene expression and subsequent expression of other transcription regulatory machinery in order to compensate. Notwithstanding these patterns, nascent gene expression in these strains indicate that the impact from the loss of each factor is highly specific.

### Antisense transcription is misregulated upon deletion of transcription regulatory factors

Loss of key transcription regulators not only affected mRNA production, but also the expression of antisense transcripts. Antisense transcripts can be classified as divergent or convergent, depending on where they occur (Mayer et al., 2015; Seila et al., 2008; Shetty et al., 2017; Xu et al., 2009). We assessed the total amount of antisense transcription from the opposite strands of protein-coding genes (**Figure 2A-B**). To determine the effects of removing transcriptional regulators on antisense transcription, we calculated the antisense:sense transcription ratios for all protein-coding genes across all deletion strains (**Figure 2C**). Interestingly, our data revealed a continuum of median antisense:sense ratios, with that of the wild-type strain near the middle of the range. The strains in which we observed the largest decrease in antisense:sense transcription ratios were those lacking factors relating to transcription elongation, such as Elf1, Spt4, and the Pol II subunit Rpb4, suggesting an asymmetry in the impact of elongation factors on sense and antisense transcripts (p = 0.03 by Fisher’s exact test). The factors whose deletions led to the largest increase in the antisense:sense transcription ratios were those involved in the regulation of histone acetylation, including members of the Rpd3S–Set2 pathway (Set2) and the major histone H4 acetyltransferase complex NuA4 (Eaf1), emphasizing the role of acetylation in antisense transcription (Carrozza et al., 2005; Churchman and Weissman, 2011; Krogan et al., 2003; Murray and Mellor, 2016; Murray et al., 2015). In many strains, changes in antisense transcription occurred in specific locations (**Figure S2A**). For example, increases in antisense transcription in the *dst1Δ* strain occurred primarily at the 3’ end; in the *set2Δ* strain, antisense transcription increased uniformly across the gene; and in the *eaf1Δ* strain, antisense transcription increased within the gene, but not at the 3’ end (**Figure 2D-F**). These findings imply that antisense transcription is a combination of different transcriptional activities regulated by separate sets of factors.

**Figure 2.**
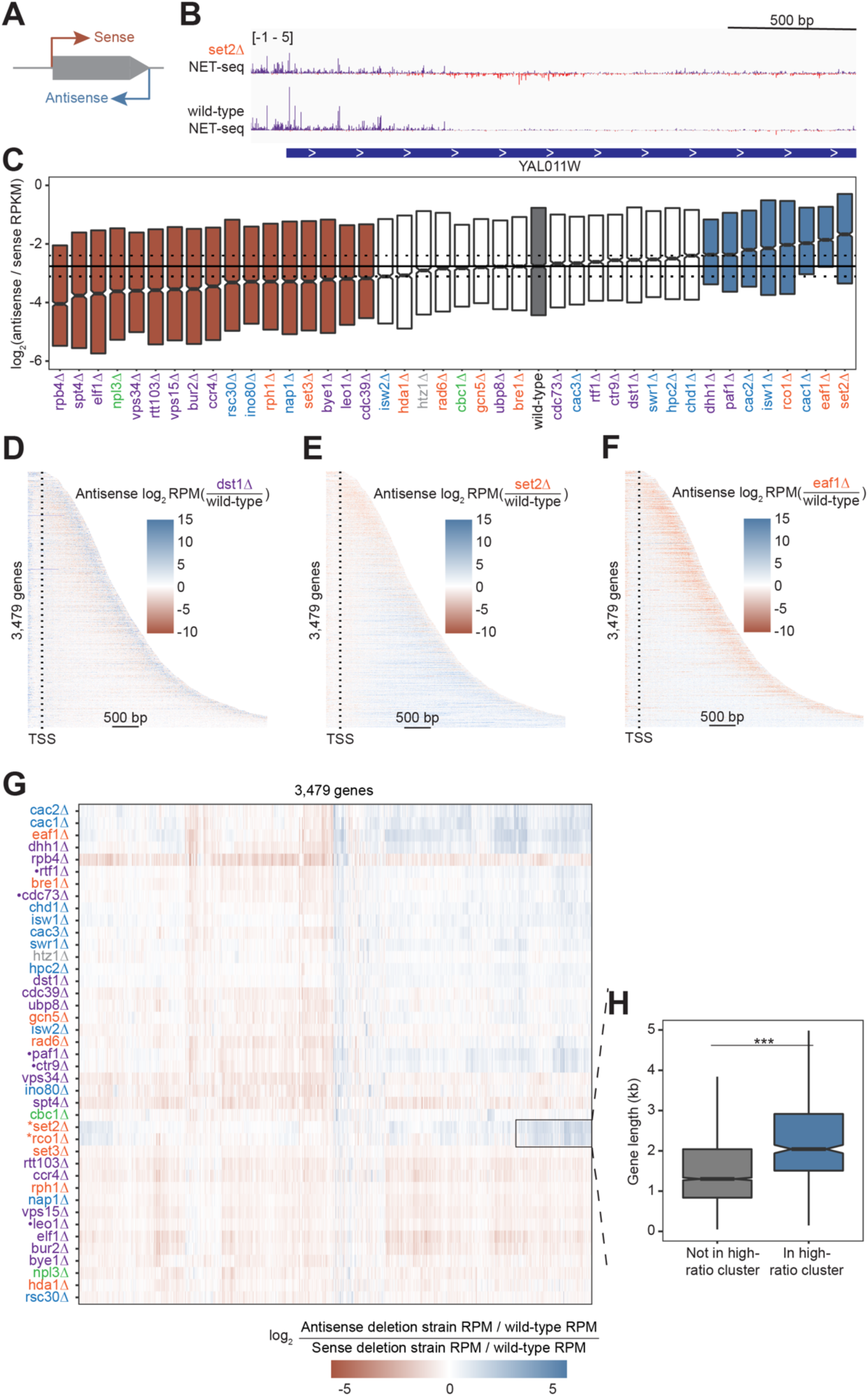
Antisense transcription is altered in most deletion strains. (A) Cartoon illustrating sense and antisense transcription of an example gene on the positive strand. (B) Wild-type and *set2Δ* NET-seq data at YAL011W. Sense and antisense are displayed in purple and red respectively. (C) Antisense:sense transcription ratios for each deletion strain compared to wild-type reveals that some strains have dramatically increased antisense transcription (*blue*) while others have much less than wild-type (*red*). Horizontal dotted line indicates the 45^th^ and 55^th^ percentile of wild-type values. (D) Heatmap of fold change in antisense transcription in the *dst1Δ* strain compared to wild-type reveals that most antisense transcription in the *dst1Δ* strain originates from the 3’ end of genes. (E-F) Same as in (D), for *set2Δ* and *eaf1Δ*, respectively. (G) Heatmap for all non-overlapping protein-coding genes, colored by the ratio of antisense to sense transcription in each deletion strain normalized to expression in wild-type. Both axes are ordered via hierarchical clustering. Box highlights a cluster of genes with high values for *set2Δ* and *rco1Δ* strains. • indicates subunits of the Paf1 complex, * indicates members of the Rpd3S-Set2 pathway (G) Boxplot illustrating the difference in distribution of gene lengths for those genes in the *set2Δ* and *rco1Δ* gene cluster. Significance was determined using a Student’s t-test (p < 0.001). Outliers are omitted from visualization.

### Losses of individual complex subunits have different effects on antisense transcription

Impacts of deletions on antisense transcription varied across the genome, i.e., not all genes experienced the same change in antisense transcription upon the loss of a given factor (**Figure 2G**). Interestingly, loss of the members of the Paf1 complex, which plays many roles in regulation of both transcription and chromatin, including the recruitment of Set2 (Chu et al., 2007; Krogan et al., 2003; Mueller and Jaehning, 2002; Schaft et al., 2003; Shi et al., 1996, 1997; Squazzo et al., 2002; Tomson and Arndt, 2013; Wade et al., 1996), did not impact antisense transcription in same genes (**Figure 2G**). Rather, clustering analysis revealed that *paf1Δ*, which had higher antisense:sense ratios, clustered with *ctr9Δ*, distant from *leo1Δ*, which has lower antisense:sense ratios, as well as from *rtf1Δ* and *cdc73Δ*, which clustered together and had closer to wild-type antisense:sense ratios.

In contrast to the Paf1 complex, deletions of two genes involved in the Rpd3S–Set2 pathway, *SET2* and *RCO1*, had similar effects, increasing antisense transcription of the same set of genes. As reported previously, these genes are longer than average (Li et al., 2007; Lickwar et al., 2009) (**Figure 2H**). However, gene length cannot fully explain why these genes had high antisense transcription, as the correlations between gene length and normalized antisense:sense transcription ratio in both *set2Δ* and *rco1Δ* strains were weak (R^2^ = 0.07 and 0.03 for *set2Δ* and *rco1Δ*, respectively; **Figure S2B**). This effect was largely specific to the *set2Δ* and *rco1Δ* strains. Most strains showed no length dependence of antisense transcription, with a few modest exceptions (*cac2Δ, dhh1Δ, eaf1Δ, paf1Δ, rtf1Δ*; top four shown in **Figure S2B**). Together, these results indicate that antisense transcription is regulated by many factors and that levels in wild-type yeast are precisely tuned by their opposing actions.

### Peaks in Pol II density across the gene body are altered in the absence of key transcription regulators

We found that Pol II density increases at loci critical for gene regulation, namely the TSS, poly(A) sites, and splice sites (SS) (F**igure S3A-D**). At the 5’ ends of genes, loss of Dst1, a homolog of the general transcription elongation factor TFIIS, dramatically increased Pol II pausing just downstream of the TSS (**Figure 3A**). We also observed peaks in Pol II density at the start of antisense transcripts opposite the 3’ ends of genes. Interestingly, deletion of *DST1* had an effect on antisense transcription similar to its impact on sense transcription (**Figure 3B, S3B**).

**Figure 3.**
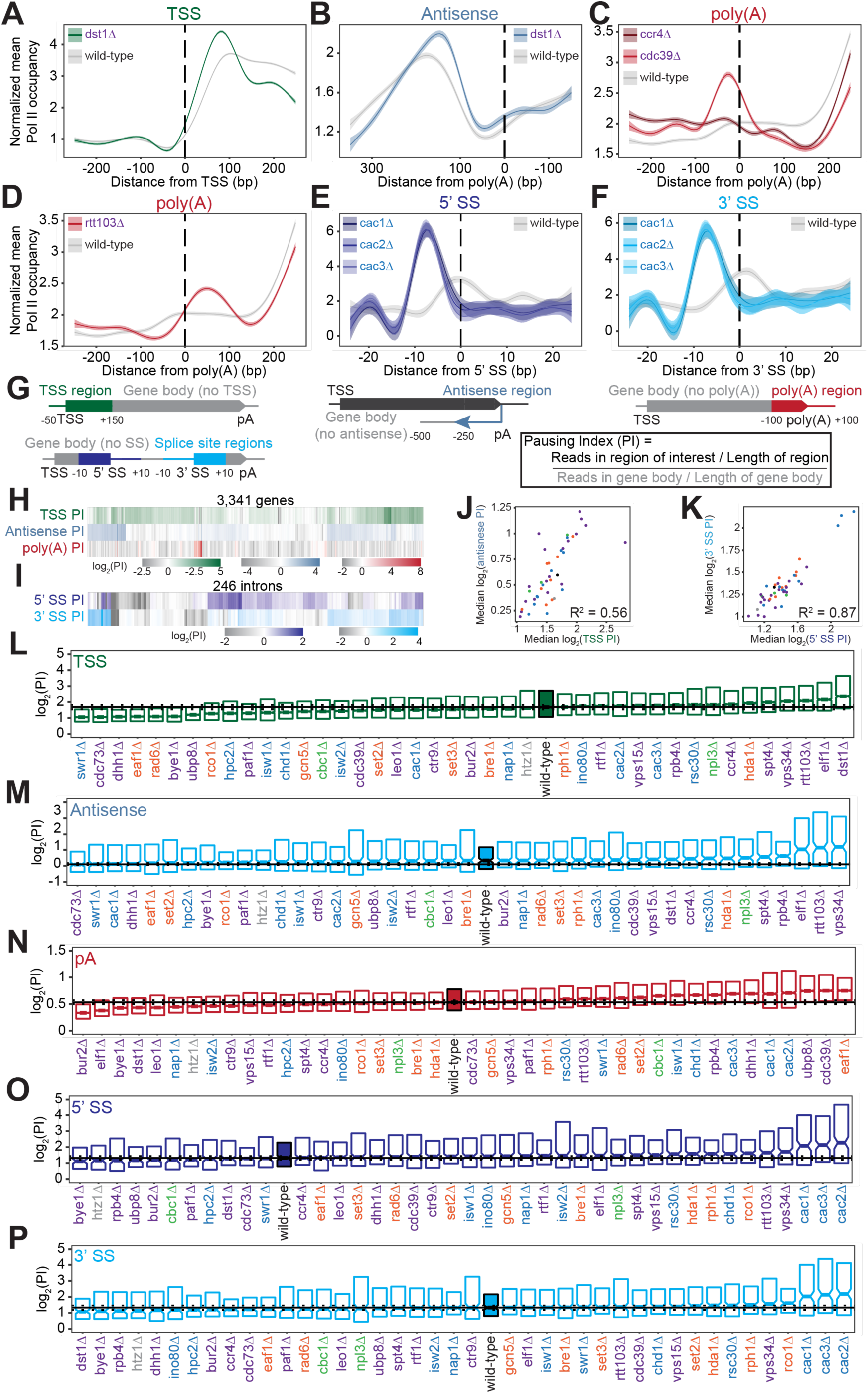
Pol II density is increased around transcription start sites, polyadenylation sites, and splice sites. (A) Metagene plot of normalized mean Pol II occupancy and the surrounding 95% confidence interval for the 500 bp surrounding the most abundant annotated transcription start sites (Pelechano et al. 2013) (n = 2,415 genes). Metagene for *dst1Δ* (*green*) can be compared to the Pol II density in the wild-type strain (*grey*). (B) Normalized mean Pol II occupancy and the surrounding 95% confidence interval for the 600 bp surrounding the most abundant annotated poly(A) sites (Pelechano et al. 2013) in the antisense orientation. Metagene for *dst1Δ* (*blue*) can be compared to the Pol II density in the wild-type strain (*grey*). (C) Normalized mean Pol II occupancy and the surrounding 95% confidence interval for the 500 bp surrounding the most abundant annotated poly(A) sites (Pelechano et al. 2013). Metagenes for subunits of the Ccr4-NOT complex deleted (*red*) can be compared to the Pol II density in the wild-type strain (*grey*). (D) Same as (C), for *rtt103Δ*. (E-F) Normalized mean Pol II occupancy and the surrounding 95% confidence interval for the 50 bp surrounding annotated 5’ and 3’ splice sites. Metagenes for subunits of the Caf1 complex deleted (*blue*) can be compared to the Pol II density in the wild-type strain (*grey*). (G) Cartoon and equation illustrating pausing index calculation. (H) Pausing index for the transcription start site (*green*), poly(A) (*red*), and 3’ antisense (*blue*) regions across genes. Horizontal axis is hierarchically clustered, revealing TSS, poly(A), and antisense pausing indices for genes in wild-type yeast. (I) Same as (H), for 5’ and 3’ splice site pausing indices. (J) Scatter plot of the median pausing indices in the TSS and 3’ antisense regions for all deletion strains. Relationship was quantified using Pearson correlation. (K) Same as in (J), comparing pausing the 5’ and 3’ splice sites surrounding introns. (L) Boxplot of TSS pausing index distributions in each deletion strain, ordered by median PI. Horizontal solid line indicates median value for wild-type yeast; dotted lines indicate the 45^th^ and 55^th^ percentile of wild-type PI values. (M-P) Same as (L), for 3’ antisense PI, poly(A) site PI, 5’ splice site PI, and 3’ splice site PI.

At the 3’ ends of genes, we observed changes in Pol II density upon loss of factors that regulate termination or polyadenylation. The screen included two subunits of the Ccr4-Not complex, which plays many roles in gene regulation including deadenylation (**Figure 3C**) (Funakoshi et al., 2007; Raisch et al., 2019; Temme et al., 2014; Tucker et al., 2002; Wahle and Winkler, 2013; Yamashita et al., 2005; Yi et al., 2018). Deletion of the scaffolding Cdc39 subunit of the complex resulted in substantial pausing before poly(A) sites, followed by reduced Pol II density. By contrast, loss of the catalytic Ccr4 subunit decreased density only downstream, with a much less prominent upstream pause (**Figure 3C**). Loss of proteins more directly involved in transcription termination, such as Rtt103, resulted in Pol II stalling just downstream of poly(A) sites, suggesting that Pol II may slow down during recruitment of this termination factor (**Figure 3D**). In these deletion strains and others, the locations of 3’-end Pol II peaks varied, with some strains exhibiting a Pol II peak before poly(A) sites and others exhibiting a peak after (**Figure S3C**), indicating that Pol II is controlled both before and after poly(A) sites.

Pol II density increases around splice sites upon the loss of several transcription regulators. For example, pause indices increased most strongly when any of the CAF-I complex components (i.e., Cac1, Cac2, Cac3) were deleted (**Figure 3E-F, S3D**). CAF-I promotes histone H3 and H4 deposition onto newly synthesized DNA (Kaufman et al., 1997), and to the best of our knowledge has not been implicated in splicing. To determine whether splicing is altered upon loss of CAF-1, we analyzed *cac2Δ* RNA-seq data (Hewawasam et al., 2018). We detected a modest but statistically significant increase in splicing in the *cac2Δ* strain relative to the wild-type (p = 0.02; **Figure S3E-F**). Thus, CAF-1 decreases Pol II density at splice sites and regulates splicing, suggesting that the complex links Pol II pausing with splicing efficiency.

To quantify Pol II pausing at each site, we defined a pausing index (PI), a length-normalized metric comparing Pol II density in the region of interest to that in the rest of the gene body (**Figure 3G**). Interestingly, genes with a high pausing index in one location did not tend to have a high index for other locations **(Figure 3H-I).** Overall, at the per gene level, there was a poor correlation between all pausing indices in the wild-type strain (e.g., TSS PI versus poly(A) PI for each gene has R^2^ = 0.06; all R^2^ ≤ 0.10, p > 0.05; **Figure S4A**). Even across each intron, pause indices differ at 5’- and 3’-splice sites although strong pausing occurs at 5’ splice sites as often as at 3’ splice sites (**Figure S4B**). Thus, pause indices vary across each gene, from the TSS to poly(A) sites, suggesting that each region of high Pol II density is regulated in a different manner.

Across deletion strains, the median pausing index varied, with the wild-type indices lying near the middle of the dynamic range (**Figure 3L-P, S4D-H).** For example, the median TSS pausing index ranged from 1.06 in *cdc73Δ* to 2.81 in *dst1Δ,* with wild-type at 1.68 (**Figure 3L**, **S3A**). The levels of antisense pausing also vary substantially across the strains (**Figure 3M**). Interestingly, the strains with the highest antisense PI had the lowest antisense:sense ratios, and vice versa (R^2^ = 0.59, p < 0.001**; Figure S2C-D**). This result implies that strong antisense pausing suppresses antisense transcription, perhaps by promoting termination and thereby preventing antisense transcription deep into gene bodies.

We asked whether the same factors are implicated in regulating the different Pol II peaks. Indeed, there was a relatively strong correlation between median TSS pausing indices and antisense pausing indices across the deletion strains (R^2^ = 0.56, p < 0.001; **Figure 3J**). Of the 10 strains with the highest TSS pausing indices, 9 were also in the top 10 for median antisense pausing indices (**Figure 3L-M**). In addition, factors that modulate pausing at splice sites tended to do so at both sites overall, but not at the same intron (R^2^ = 0.87, p < 0.001; **Figure 3K, S4B**). However, we did not observe similar relationships between other pause indices (**Figure S4C**). Thus, similar regulatory mechanisms function at both 5’ and 3’ splice sites and at both sense and antisense transcription start sites.

### Pol II pausing propensity and location are affected by deletion of transcription regulators

Along with identifying regions of elevated Pol II density, NET-seq data pinpoints the precise positions that Pol II pauses at within regions of high Pol II density and elsewhere. These precise sites of Pol II pausing at single nucleotides are reminisant of *in vitro* RNA polymerase pausing observed at specific positions of DNA templates (Galburt et al., 2007; Hodges et al., 2009; Kingston and Chamberlin, 1981; Mayer et al., 2017; Wang et al., 1998). We systematically identified pause sites in strains with sufficient coverage as positions with read densities that deviate from the statistical fluctuations of the surrounding 200 nucleotides, modeled as a negative binomial distribution (>3 standard deviations from the mean; **Figure 4A-B**). We found thousands of reproducible pauses at single nucleotides in highly expressed genes across all deletion strains (irreproducible discovery rate <1%; **Figure S5A-B**) (Li et al., 2011).

**Figure 4.**
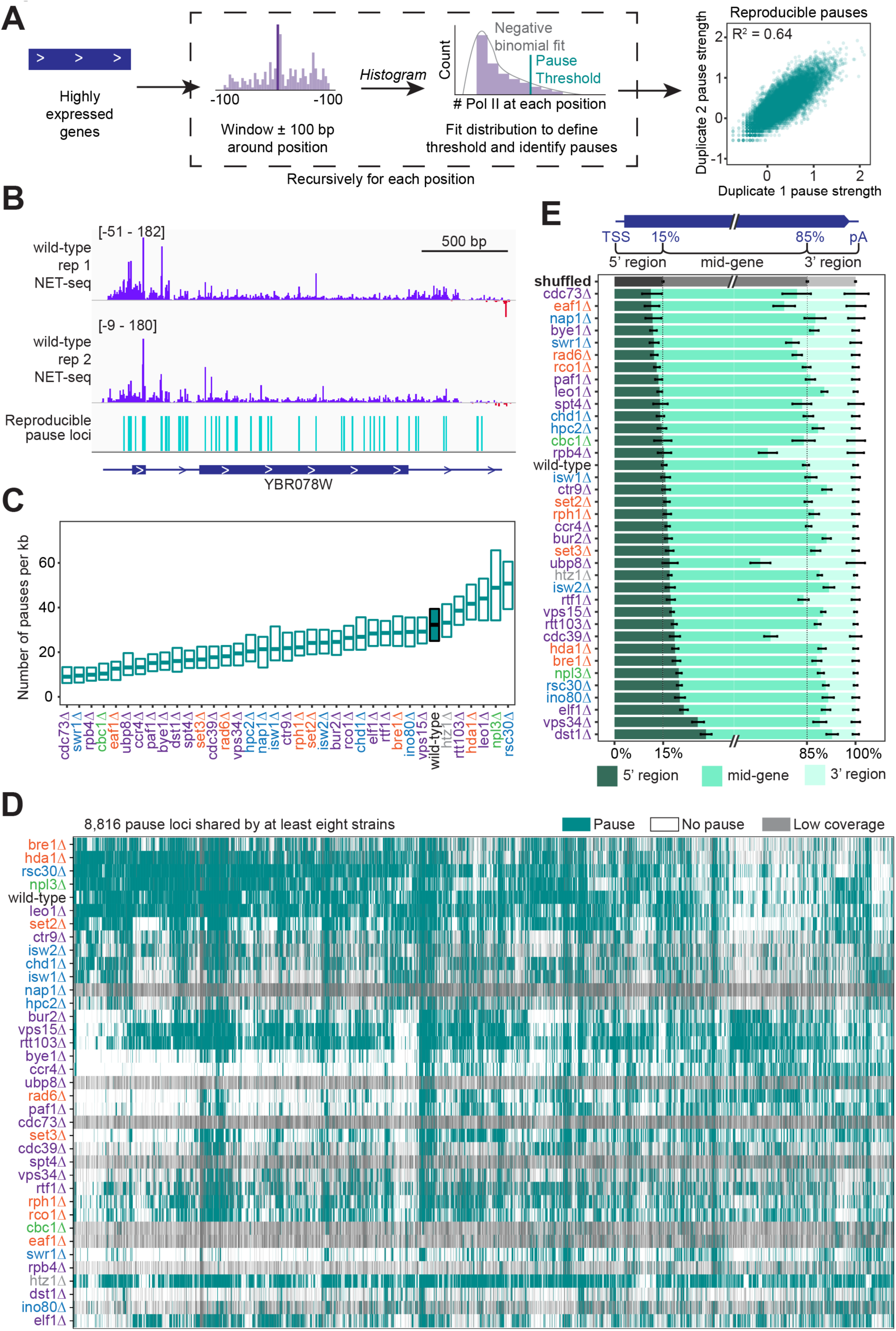
Trends in Pol II pausing behavior at single-nucleotide resolution across deletion strains. (A) Cartoon illustrating algorithm for robust and reproducible Pol II pause detection. Scatter plot shows correlation between pause strength in two different wild-type replicates. Pause strength is measured by log_10_(RPM) at pause loci, and only reproducible pauses are shown (IDR ≥ 1%). (B) Example of Pol II density on the positive (*purple*) and negative (*red*) strands, as measured by NET-seq in two wild-type replicates. Pauses that meet the 1% IDR reproducibility threshold are shown as blue vertical lines. (C) Boxplot of the distribution of Pol II pause densities, the number of pauses per kilobase examined, in each deletion strain, ordered by median pausing density. Whiskers and outliers were removed for visualization. (D) Hierarchically clustered heatmap of 8816 Pol II pause loci across the genome reveals locations of pauses shared by multiple deletion strains. Heatmap is colored based on if that locus was identified as a pause (*teal*), not a pause (*white*), or if there was not sufficient coverage to determine pause status (*grey*). Analyses conducted only on deletion strains with biological replicates and only at loci at which there was enough coverage to determine the absence of a Pol II pause in at least one deletion strain. (E) The percent of Pol II pause loci located in the 5’ gene region, mid-gene, and 3’ gene region varies across deletion strains. 5’ gene region was identified for each well-expressed gene as extending from the transcription start site to the 15^th^ percentile of the gene length. Similarly, the 3’ gene region was defined as the last 15^th^ percentile of the gene length, with the mid-gene region spanning in between. The control (*grey*) was created by scrambling all identified pauses across all deletion strains within the genes they were identified in. Rows are ordered by the percent of pauses found in the 5’ region.

Pause site density, or the number of pauses per kilobase, varied widely across deletion strains (**Figure 4C**) that cannot be explained by differences in sequencing depth across deletion strains (R^2^ = 0.001, p =0.845; **Figure S5C**). In the wild-type strain, we found Pol II pause sites on average every 31 bp. Loss of most transcription elongation factors resulted in a lower pause site density relative to the wild-type (**Figure 4C**).

However, some of the deletion strains exhibited more pausing overall; upon loss of capping factor Npl3, for example, 35% of all NET-seq reads mapping to highly expressed genes constituted pause sites, versus only 21% in the wild-type (**Figure S5B**). The increased Pol II pause site density in *npl3Δ* cells is consistent with its role in stimulating Pol II elongation (**Figure 4C**) (Dermody et al., 2008).

The pause loci for each strain included many that were not observed in wild-type yeast (**Figures 4D, S5D**). Indeed, when the sets of pause loci are used to cluster deletion strains by principal component analysis, the wild-type strain stands away from most strains (**Figure S5D**). However, some deletion strains shared many pause sites with those observed under in the wild-type: 81% of pause sites identified in wild-type yeast were also identified in the *htz1Δ* strain, consistent with its confined role at the +1 nucleosome (Bagchi et al., 2020; Zhang et al., 2005).

We wondered whether loss of related factors would lead to the same sets of pause sites. We first identified all pauses observed in at least 8 strains and used the presence or absence of these pauses in each strain to perform hierarchical clustering (**Figure 4D**). *dst1Δ* pause sites clustered far away from those in wild-type cells, consistent with the backtracking role of Dst1 that leads to downstream-shifted pause sites (Churchman and Weissman, 2011; Noe Gonzalez et al., 2021). H2B ubiquitidation increases the nucleosomal barrier to Pol II (Chen et al., 2019), so alterations to histone ubiquitination might lead to new pause sites. Interestingly, pause sites after the loss of Rad6, Ubp8, Paf1 and Cdc73 all cluster together. Rad6 and Ubp8 ubiquitinates and deubiquitinates H2B respectively (Amerik et al., 2000; Jentsch et al., 1987). Paf1 and Cdc73, members of the Paf1 complex, are responsible for recruiting Rad6 to chromatin (Kim and Roeder, 2009). The clustering of these factors indicates a role for H2B ubiquidation in determining the locations of many pause sites. Finally, we figured that differences in nucleosome positioning may lead to differential pause sites usage, so we inspected how pause sites change after the loss of different chromatin remodelers. Interestingly, we observed that loss of ISWI and CHD chromatin remodelers Isw1, Isw2, and Chd1 lead to pause sites that cluster together (**Figure 4D**). For example, most of the pause sites observed in *isw1Δ* (76%) were also observed in *chd1Δ*, consistent with their joint roles in maintaining chromatin structure (Smolle et al., 2012). In contrast, loss of Ino80, Rsc30 and Swr1 all lead to distinct sets of pause sites (**Figure 4D**). *isw1Δ* pause sites cluster as far from those of wild-type as *dst1Δ* pause sites, perhaps due to the large-scale disruption of chromatin structure after Ino80 loss (Kubik et al., 2019).

Pol II pause sites in the wild-type strain were distributed evenly throughout gene bodies (**Figure 4E**). By contrast, deletion strains exhibited a range between two-fold decreased to two-fold increased Pol II pause sites in the 3’ regions of genes, with slightly less variability at the 5’ regions of genes relative to a scrambled control or wild-type pausing (**Figure 4E**). The enrichment of pause sites at 5’ end and 3’ regions generally correspond with our pausing index results (**Figure 3H, L, N**). For example, deletion of *DST1* approximately doubled pause loci in the 5’ regions at the expense of pausing in 3’ regions. However, in general, changes in 5’ vs 3’ pause sites in deletion strains were not correlated (**Figure 4E**). We find substantially more pause sites at the 3’ regions of genes in *rpb4Δ*. Rpb4 is a Pol II subunit that dissociates with the complex at the ends of genes (Mosley et al., 2013) and is responsible for sustained transcription elongation through the 3’ ends of genes (Runner et al., 2008). Thus, Rpb4 prevents Pol II from pausing at the 3’ regions of genes that may protect from premature termination before the canonical 3’ cleavage site is transcribed. Similarly, more 3’ pause sites are found in the *ubp8Δ* strain, consistent with the global increase in this strain of H2B ubiquidylation, a mark that increases the nucleosomal barrier to Pol II and is coincident with Pol II pausing at transcription termination sites (Bonnet et al., 2014; Chen et al., 2019; Harlen et al., 2016). Together, these data show how the chromatin landscape and transcriptional regulatory network of the cell dictate sites of Pol II pausing that in turn controls where and for how long Pol II pauses during elongation.

### Chromatin features can accurately predict Pol II pausing locations in deletions strains

Given the number of reproducible pause sites we identified, we next investigated whether we could determine which genomic features, if any, were responsible for the pause sites. *In vitro* studies have shown that Pol II pausing has many causes, including specific DNA sequences, nucleosomes, and histone modifications (Bintu et al., 2012; Herbert et al., 2006; Hodges et al., 2009; Kassavetis and Chamberlin, 1981; Kireeva and Kashlev, 2009; Kireeva et al., 2005; Shaevitz et al., 2003). *In vivo*, the dominant factors globally associated with Pol II pause sites remain unclear, although sequence elements, transcription factors, nucleosomes, and CTD modifications have all been connected to Pol II pausing (Alexander et al., 2010; Churchman and Weissman, 2011; Gajos et al., 2021; Nechaev et al., 2010; Noe Gonzalez et al., 2021; Nojima et al., 2018; Shukla et al., 2011). Recently, DNA sequence and shape were shown to be important contributors to pause site locations in human cells (Gajos et al., 2021). We first asked whether specific DNA sequences were connected with Pol II pausing loci. Previous studies reported that Pol II has a strong bias toward pausing at adenine (Churchman and Weissman, 2011), which we also observed here. More specifically, we observed a 3.4-fold enrichment of real Pol II pause sites at T<u>A</u>T trinucleotide sequences relative to shuffled control sites in the same well-expressed genes (**Figure 5A**). The shape of the DNA itself, as predicted from sequence, also appears to inform the location and propensity for Pol II to stall: DNA low helix twist values were more common under real pause loci than in the shuffled control (**Figure 5B**). These observations were consistent, as the AT dinucleotide step has a low average twist angle of 32.1° (Ussery, 2002).

**Figure 5.**
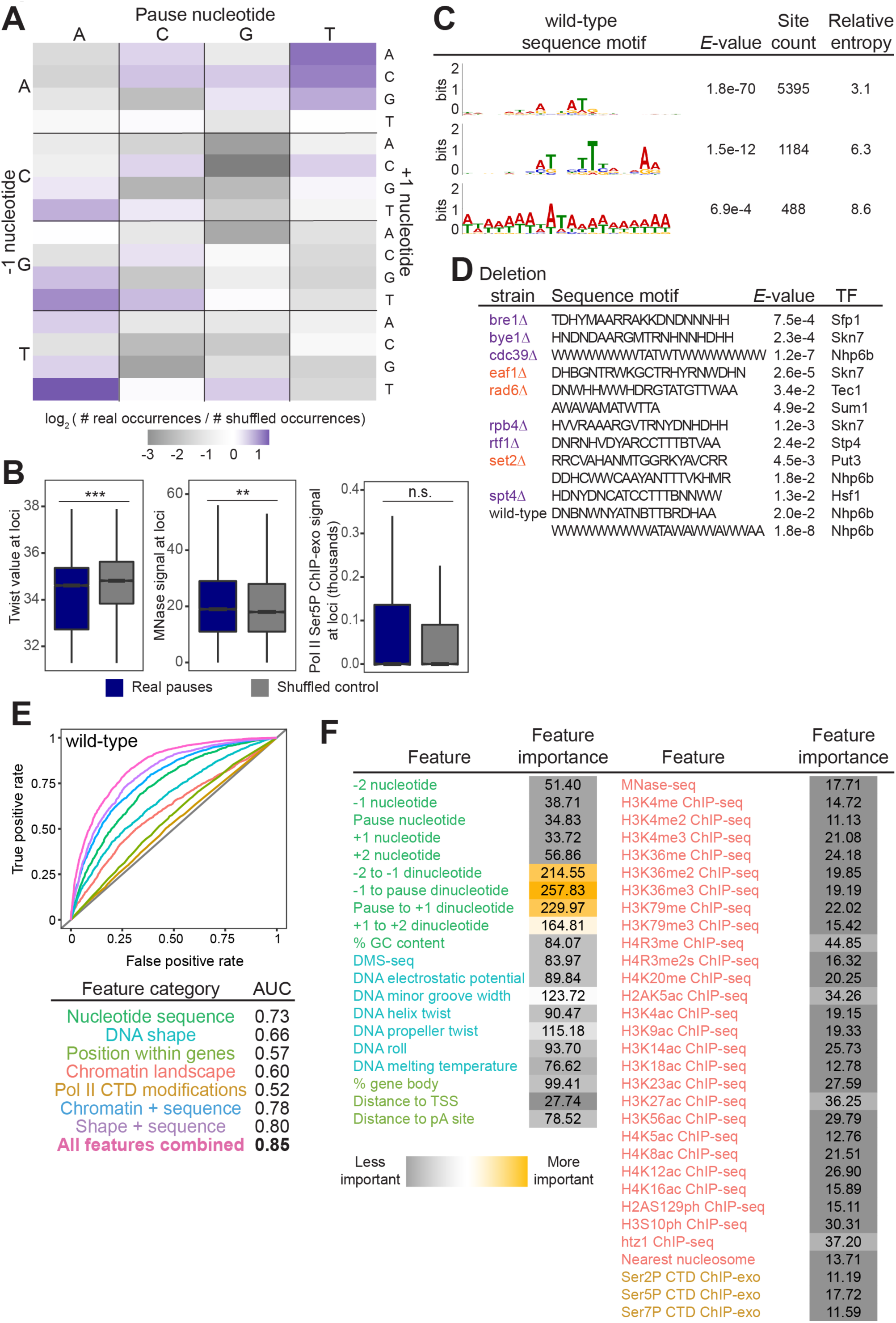
Chromatin features explain the location of some Pol II pauses in wild-type yeast. (A) Heatmap illustrating the relative frequency of each trinucleotide sequence surrounding real and shuffled control pauses centered on Pol II pauses identified in wild-type. (B, left) Comparison in the distribution of values for twist values underlying Pol II pauses in wild-type yeast (n = 13,994) compared to a shuffled control, in which the same number of pauses are shuffled, maintaining the same number of pauses within each well-expressed gene. Differences between the real and shuffled distributions were determined as significantly significant by a Student’s t-test with Bonferroni correction for multiple hypotheses. P-values are reported in Table S5. (* adjusted p-value ≤ 0.05; ** adjusted p-value ≤ 0.01; *** adjusted p-value ≤ 0.001). Also shown for MNase-seq signal (center) and Ser5P CTD ChIP-exo signal (right).(C) Table showing the three significant motifs identified under Pol II pauses in the wild-type strain. All analysis was performed using the MEME suite of tools. Significant motifs were those with an *E*-value greater than 0.05. Pause sites were scrambled within well-expressed genes to be used as a negative control and to calculate enrichment of motifs. (D) Table with all sequence motifs underlying pauses across deletion strains that are significantly similar to known transcription factor binding motifs. Only the top match, as assessed by E-value, is reported. (E) Receiver operating characteristic (ROC) curve from a random forest classifier that measures the predictive value of chromatin features on Pol II pauses in wild-type yeast (10,495 training and 3,499 training loci). (F) Table of all features used in random forest classifier for pause loci classification and the importance of each feature. Feature importance is calculated as the mean decrease in accuracy upon removing that feature from the model.

Beyond the trinucleotides, significantly enriched sequence motifs were also associated with Pol II pause sites in most deletion strains (**Table S4**), including three motifs related to pauses in the wild-type strain (**Figure 5C**). Notably, not all motifs are shared across strains, and upon deletion of some factors, new motifs were associated with Pol II pause sites. 13 of the 26 identified sequence motifs with high relative entropies significantly matched known transcription factor binding site motifs (**Figure 5D**). Thus, it is likely that Pol II pause sites can partially, but not fully, be explained by DNA sequence and/or proteins binding to DNA.

In addition to the structure of the DNA itself, chromatin features, such as nucleosome positions and histone modifications, are also connected to Pol II pausing behavior. To search broadly for genomic features underlying sites of Pol II pausing, we evaluated 51 chromatin features (**Table S5**), including nucleotide sequence, DNA shape, position of pauses within a gene, histone modifications, and Pol II CTD phosphorylation marks. 35 out of 42 exhibited a statistically significant difference between real wild-type pause sites and shuffled controls (the remaining nine of the 51 are sequence features that cannot be compared on a numeric scale) (**Figure S6A, Table S6**). For example, the MNase-seq signal around pause loci and the distance to the nearest nucleosome differed significantly between real and shuffled pause sites (**Figure 5B, S6A**), consistent with observations of pauses at nucleosomes (Churchman and Weissman, 2011). Interestingly, Ser2, Ser5 and Ser7 phosphorylation of the Pol II CTD did not differ relative to random positions, indicating that connections between Pol II phosphorylation and pausing at intron-exon boundaries is specific to pausing at those loci (Alexander et al., 2010). Among the features that differed significantly was DNA melting temperature, which was previously shown to influence Pol II stalling (Nechaev et al., 2010).

To determine whether any features could predict where Pol II pauses, we created a random forest classifier to discriminate between real and shuffled control Pol II pause sites based on the surrounding chromatin features. A random forest classifier using all 51 features performed well (AUC = 0.85, **Figure 5E**) relative to a random model (AUC = 0.5) at classifying Pol II pauses in wild-type yeast. The most critical features for accurate identification of Pol II pause sites were DNA sequence surrounding the pause locus and topology features of the DNA at that locus (**Figure 5F**). Together, these analyses showed that DNA sequence and shape contribute strongly to Pol II pause locations, but their effects are enhanced by many other chromatin features.

To ask whether prediction models for Pol II pausing vary in different regulatory and chromatin landscapes, we built random forest models for each deletion strain. Across all deletion strains, an AUC of at least 0.78 was attained. These AUC values were only partially correlated with the total number of pauses detected in each deletion strain (R^2^ = 0.37, p =0.000064; **Figure S7A**). Although nucleotide sequence and DNA shape were the most important features for classifying Pol II pause loci in the wild-type and many deletion strains, models for a subset of strains (including *cdc39Δ, dst1Δ, ubp8Δ*) revealed that wild-type chromatin modifications were more powerful for Pol II classification (**Figure 6A, S7B-E**). We next performed a transfer of learning analysis to ask how each model would perform when predicting pauses in other strains. When trained on Pol II pause sites identified in wild-type yeast, the AUC when testing on pauses across all other strains ranged from 0.53 (*cbc1Δ*) to 0.82 (*vps15Δ*), revealing the differences across the strains (**Figure 6B**). We previously observed that loss of Dst1 leads to ∼75% of pause sites to shift downstream(Churchman and Weissman, 2011). Thus, training a model on *dst1Δ* pause sites should not do well to predict pauses in another strain. Indeed, a model trained on *dst1Δ* pause sites performed well in predicting *dst1Δ* pause sites (AUC = 0.83), however it performed the worst of all models in predicting pause sites in other deletion strains, obtaining a median AUC of 0.63 across them. These models indicate that the nucleotide sequence, DNA topology, position within a gene, and chromatin landscape all play roles in determining the location of Pol II pauses during transcription elongation.

**Figure 6.**
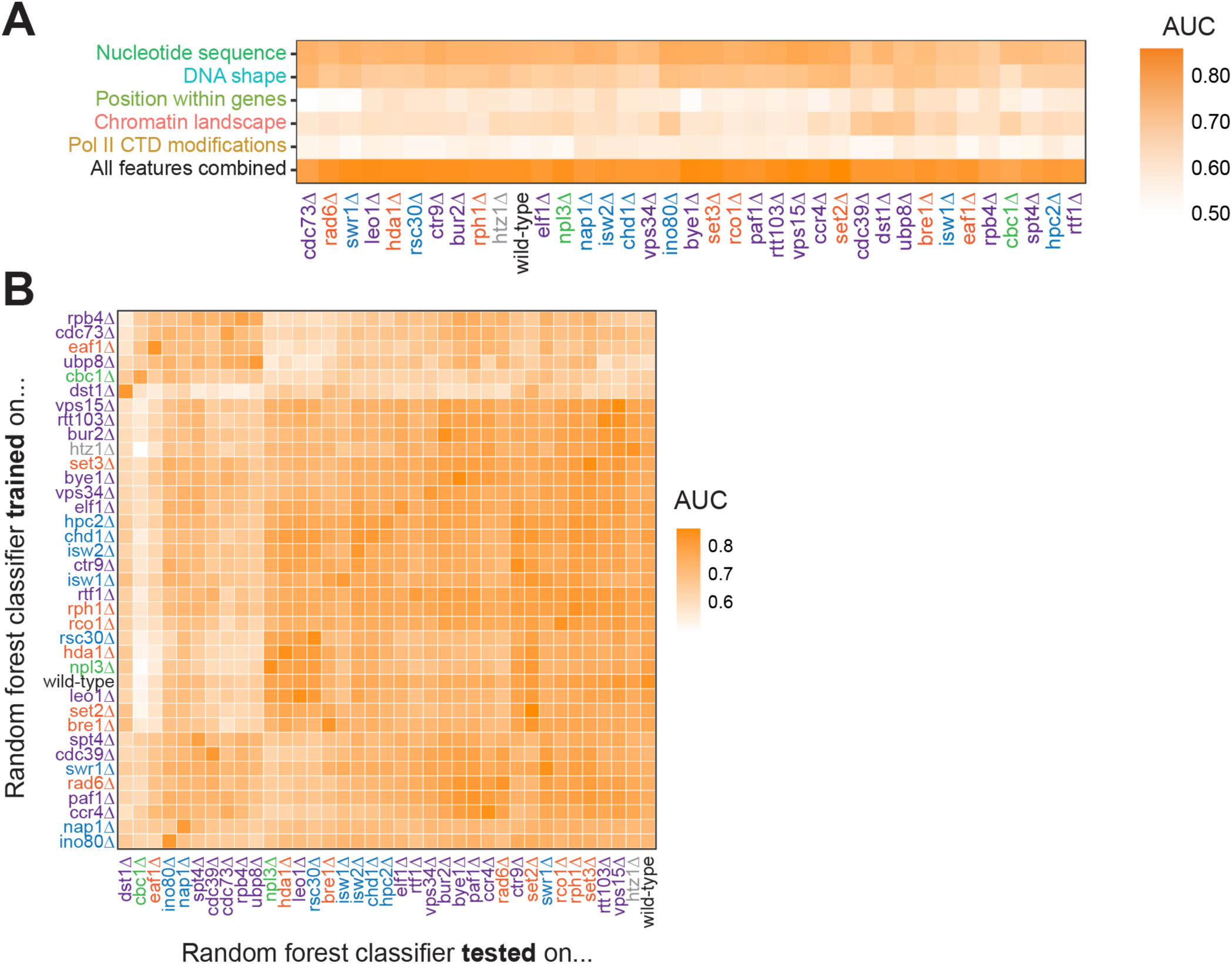
Random forest classifiers can predict Pol II pause loci across deletion strains, with different feature importance values across deletion strains. (A) Heatmap illustrating the mean AUC for the random forest classifier when trained (75% of loci) and tested (25% of loci) on each deletion strain. Deletion strains are hierarchically clustered along the x-axis. (B) Heatmap showing the AUC values from random forest classifiers trained on all pauses from one deletion strain (y-axis) and tested on those unique pauses observed in another deletion strain (x-axis). Both axes are hierarchically clustered to reveal similarities in AUC values across deletion strains. Tiles when the same training and testing strain are indicated are colored according to the AUC for that deletion strain when 75% of pauses in that deletion strain are used for training and the remaining 25% are used for testing as reported in (A).

### Pol II pausing and antisense transcription are linked to overall gene transcription

Antisense transcription can repress sense transcription through transcriptional interference (Lenstra et al., 2015; Nevers et al., 2018; Scruggs et al., 2015), and strong Pol II pausing can lead to low transcriptional output or early termination (Gressel et al., 2017; Shao and Zeitlinger, 2017; Steurer et al., 2018). However, it remains unclear whether changes in transcriptional output are generally connected to changes in Pol II pausing or antisense transcription. Through a meta-analysis of the NET-seq data sets, we compared changes in gene transcription to changes in Pol II pausing and antisense transcription across a range of transcription regulatory landscapes.

First, we compared levels of sense and antisense transcription in all strains analyzed and saw a modest correlation between fold change in the antisense:sense ratios and fold change in expression (r = 0.38) (**Figure 7A**). For the TSS pausing index, we do not observe a correlation between changes in expression and changes in pausing (r =0.062 for all deletion strains) (**Figure 7A**). We reasoned that genes exhibiting sensitivity to alterations in the gene regulatory landscape might exhibit greater sensitivity to changes in pausing or antisense transcription. Indeed, for frequently-regulated genes (differentially transcribed in eight or more strains), correlations between antisense:sense ratios and fold change in expression ( r = 0.6) or TSS pausing index and fold change in expression (r = 0.13) increase (**Figure 7B**).

**Figure 7.**
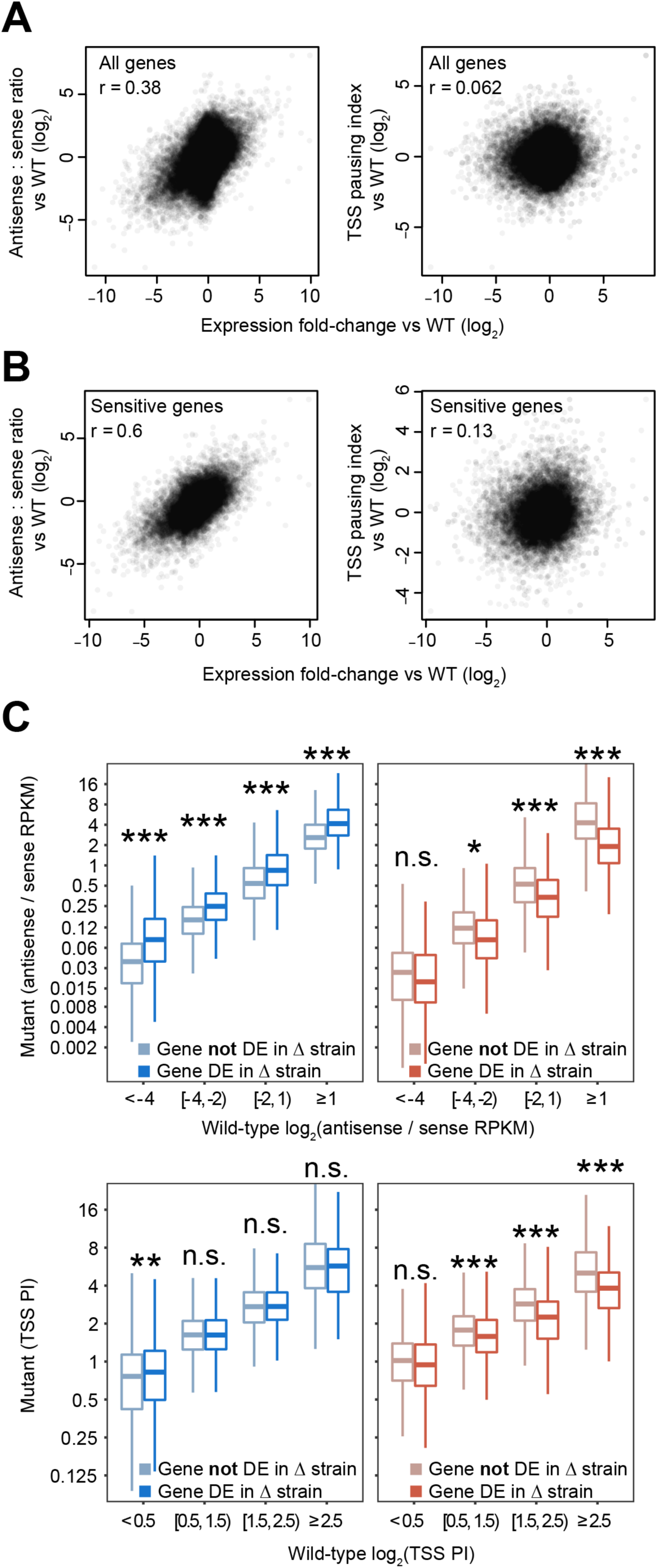
Antisense transcription and Pol II pausing around the TSS are intimately linked to gene expression across deletion strains. (A) Scatter plots of antisense:sense ratio fold-change compared to wild type (*left*) and TSS pausing index fold-change compared to wild type (*right*) vs expression fold-change compared to wild type for all genes across all strains. Pearson r value is shown. (B) Scatter plots of antisense:sense ratio fold-change compared to wild type (*left*) and TSS pausing index fold-change compared to wild type (*right*) vs expression fold-change compared to wild type for sensitive genes across all strains. Sensitive genes are those that were differentially expressed in at least eight deletion strains. Pearson r value is shown. (C) Boxplots illustrating distributions of antisense:sense transcription ratios (top panels) or TSS pausing index (bottom panels) in deletion strains for binned antisense:sense ratios in wild-type. The only genes plotted were those that were differentially up-regulated (*left*) or down-regulated (right) in at least eight deletion strains. Distributions in each bin were split by whether that gene-deletion strain combination was differentially transcribed (n = 239 genes up-regulated in at least 8 deletion strains, n = 316 genes down-regulated in at least 8 deletion strains). Asterisks indicate p-value (* < 0.05, ** < 0.01, *** < 0.001, n.s. not significant) from a Student’s t-test comparing the distribution of values from differentially expressed and not differentially expressed genes.

To determine whether pausing and antisense transcription are linked to gene regulation, we asked how pausing and/or antisense transcription changes specifically when frequently-regulated genes are differentially expressed. As each of these genes is not differentially expressed in many deletion strains, we compared the pausing and antisense transcription of the genes in strains where the gene is regulated to strains where it is not. We found that frequently up-regulated genes exhibited higher antisense:sense ratios when they were up-regulated relative to when they were not differentially transcribed (**Figure 7C**). By contrast, frequently down-regulated genes had lower antisense:sense ratios when they were down-regulated (**Figure 7C**). With respect to pausing, genes that were commonly down-regulated exhibited decreases in their TSS pausing specifically in strains where their expression levels change (**Figure 7C**). Frequently up-regulated genes had more pausing near the TSS when they were up-regulated than when they did not experience a change in expression (**Figure 7C**), although the effect size is too low to be considered statistically significant for genes already experiencing high levels of pausing. Together, genes that are frequently down-regulated tend to exhibit a decrease in antisense transcription and TSS pausing, whereas frequently up-regulated genes tend to exhibit an increase in both, but only when regulated. Thus, there is a relationship between TSS pausing, antisense transcription and gene regulation that argue against a general repressive role for antisense transcription and Pol II pausing.

## DISCUSSION

Advances in high-throughput sequencing of nascent RNA have revealed that, in many eukaryotes, the vast majority of the genome is transcribed (Hangauer et al., 2013; Struhl, 2007). Nevertheless, this broad transcriptional activity is one of the most highly regulated processes within the cell. Multiple levels of regulation are orchestrated by DNA sequence, transcription factors, RNA processing factors, and chromatin modulators. Here, we used NET-seq to study 41 factors with connections to transcription elongation and discovered the remarkable tunability of transcription elongation. For all of the transcriptional phenotypes analyzed, the wild-type strain fell in the middle of the dynamic range observed across the deletion strains, revealing the intricate balance of multiple aspects of transcriptional activity.

The 41 factors chosen for this study were previously annotated to regulate transcription elongation. However, loss of each factor had a unique impact on gene expression, suggesting that genes are differentially sensitized to perturbations of the transcription regulatory network. Levels of antisense transcription in the deletion strains vary across a broad dynamic range, revealing that antisense transcription is finely tuned by many factors. Interestingly, loss of 17 factors decreased the antisense:sense transcription ratio in cells (**Figure 2C**), indicating that it is possible to suppress antisense transcription further than what is observed in wild-type. Conversely, loss of eight factors increased antisense:sense ratios. Together, these results imply that wild-type antisense transcription is balanced by the influence of many factors and, in turn, can be precisely controlled. The possibility of tight control of antisense transcription indicates that regulatory mechanisms may exist where antisense transcription impacts sense transcription. Indeed, differentially transcribed genes showed pronounced changes in their antisense:sense transcription ratios, especially for a subset of sensitive genes that are differentially transcribed in many of the deletion strains.

Peaks of Pol II density were detected near TSSs, poly(A) sites, and both 5’ and 3’ splice sites. Interestingly, factors that impacted pausing at the 5’ ends of genes were not the same as those that impacted pausing at 3’ ends or at splice sites. Clearly, different mechanisms regulate Pol II pausing at different points during elongation. However, pausing around the TSS and pausing during antisense transcription were controlled by a similar set of factors, suggesting the existence of a checkpoint early in transcription, in the sense and antisense directions.

Within the regions of elevated Pol II density and across the gene body are discrete pauses at single nucleotides throughout elongation. Our data indicated that in wild-type cells, the density and intensity of these pause sites are precisely balanced by numerous factors. For example, the set of Pol II positions where Pol II pauses vary substantially across the deletion strains (**Figure 4D**), indicating that there are a large number of positions where Pol II could pause in wild-type cells, but the presence of these factors modulates the pausing landscape such that they are not utilized.

We identified genes that are sensitive to alterations of the transcription elongation regulatory network and found their changes in expression are accompanied by changes in Pol II pausing. When these genes are down-regulated, Pol II peaks after TSSs are smaller and vice versa for up-regulated genes, suggesting a possible stimulatory role of these pauses (**Figure 7C**). However, this result is in contrast to our observation that higher antisense pausing was generally associated with lower levels of antisense transcription overall (**Figure S2C-D**). Thus, pausing in the antisense direction seems to lead to tighter quality control through earlier termination. While in the sense direction, it may enhance expression levels for some genes (i.e. the sensitive genes identified in this screen), perhaps by keeping the promoter region open to encourage more initiation as described in mammalian cells (Scruggs et al., 2015).

This work reveals the complex regulation of transcription elongation by a network of factors. In addition, it serves as a resource of NET-seq data to explore more specific hypothesis-driven research questions relating to individual factors and an open-source code base with which to analyze these data. Many of the transcription elongation regulators studied here are conserved in all domains of life, as are many of the transcriptional phenotypes we examined, including antisense transcription and Pol II pausing. These insights into transcription regulation in *S. cerevisiae* will serve as a foundation for learning more about transcription in multicellular eukaryotes.

## METHODS

### Yeast mutant generation

To create deletion mutants of the 41 factors analyzed, the parent strain YSC001 (BY4741 *rpb3::rpb3-3xFLAG NAT*) (Churchman and Weissman, 2011) was transformed with PCR products of the HIS3 gene flanked by 40 bp of homology upstream and downstream of the start and stop codons for the gene of interest. Standard lithium acetate transformations were used.

### NET-seq library generation

Cultures for NET-seq were prepared as described in (Churchman and Weissman, 2012). Briefly, overnight cultures from single yeast colonies grown in YPD were diluted to OD_600_ = 0.05 in 1L of YPD medium and grown at 30°C shaking at 200 rpm until reaching an OD_600_ = 0.6 - 0.8. Cultures were then filtered over 0.45mm-pore-size nitrocellulose filters (Whatman). Yeast was scraped off the filter with a spatula pre-chilled in liquid nitrogen and plunged directly into liquid nitrogen as described in (Churchman and Weissman, 2012). Mixer mill pulverization was performed using the conditions described above for 6 cycles. NET-seq growth conditions, IPs, and isolation of nascent RNA and library construction were carried out as described in (Churchman and Weissman, 2012). A random hexamer sequence was added to the linker to improve ligation efficiency and allow for the removal of any library biases generated from the RT step as described in (Mayer et al., 2015). After library construction, the size distribution of the library was determined by using a 2100 Bioanalyzer (Agilent) and library concentrations were determined by Qubit 2.0 fluorometer (Invitrogen). 3’ end sequencing of all samples was carried out on an Illumina NextSeq 500 with a read length of 75 bp. For analysis of *cac1Δ*, *cac2Δ*, and *cac3Δ*, raw Fastq files were obtained from (Marquardt et al., 2014) and re-aligned using the parameters described below.

### Processing and alignment of NET-seq data

The adapter sequence (ATCTCGTATGCCGTCTTCTGCTTG) was removed using cutadapt with the following parameters: -O 3 -m 1 --length-tag ‘length=’. Raw fastq files were filtered using PRINSEQ (http://prinseq.sourceforge.net/) with the following parameters: -no_qual_header -min_len 7 -min_qual_mean 20 -trim_right 1 -trim_ns_right 1 -trim_qual_right 20 -trim_qual_type mean -trim_qual_window 5 -trim_qual_step 1. Random hexamer linker sequences (the first 6 nucleotides at the 5’ end of the read) were removed using custom Python scripts, but remained associated with the read. Reads were then aligned to the SacCer3 genome obtained from the *Saccharomyces* Genome Database using the TopHat2 aligner (Kim et al., 2013) with the following parameters: --read-mismatches 3 --read-gap-length 2 --read-edit-dist 3 --min-anchor-length 8 --splice-mismatches 1 --min-intron-length 50 --max-intron-length 1200 --max-insertion-length 3 --max-deletion-length 3 --num-threads --max-multihits 100 --library-type fr-firststrand --segment-mismatches 3 --no-coverage-search --segment-length 20 --min-coverage-intron 50 --max-coverage-intron 100000 --min-segment-intron 50 --max-segment-intron 500000 --b2-sensitive. To avoid any biases toward favoring annotated regions, the alignment was performed without providing a transcriptome. Reverse transcription mispriming events were identified and removed where molecular barcode sequences correspond exactly to the genomic sequence adjacent to the aligned read. With NET-seq, the 5’ end of the sequencing, which corresponds to the 3’ end of the nascent RNA fragment, is recorded with a custom Python script using the HTSeq package (Anders et al., 2015). NET-seq data were normalized by million mapped reads. Replicate correlations were performed comparing RPKM of each gene in each replicate; replicates were considered highly correlated with a Pearson correlation of R^2^ ≥ 0.75. Raw NET-seq data of highly correlated replicates were merged, and then re-normalized by million mapped reads. For analysis of *rco1Δ*, raw Fastq files were obtained from (Churchman and Weissman, 2011) and re-aligned using the parameters described below.

### Differential gene transcription and gene ontology enrichment analysis

Expression levels were determined for each gene using the entire sequencing read (rather than only the 3’ end, as typical when determining Pol II location from NET-seq data) and normalized using DESeq2 (Love et al., 2014). Differential transcription analysis between deletion strains (with replicates) and wild-type strains was performed using DESeq2 (Love et al., 2014) for all UCSC known genes. Genes were considered differentially transcribed if they had an adjusted p-value < 0.05 and an absolute log_2_ fold change > 1.0. The relationship between the number of differentially transcribed genes identified and growth rate (i.e., doubling time) of strains was quantified using Pearson correlation. Because the *cac1Δ*, *cac2Δ*, and *cac3Δ* strains were constructed in a different lab (Marquardt et al., 2014), they were excluded from this analysis.

GO term enrichment analysis was performed using the PANTHER GO Enrichment Analysis (Ashburner et al., 2000; Mi et al., 2019; The Gene Ontology Consortium, 2019; Thomas et al., 2003) and enriched terms for biological processes, molecular function, and cellular components were recorded for up-regulated differentially transcribed genes, down-regulated differentially transcribed genes, and all differentially transcribed genes. Fold enrichment and false discovery rates for each GO by deletion strain pair are reported in (**Table S3**).

### Trends in differentially transcribed genes

Frequently regulated genes were identified as those that had been differentially transcribed in 8 or more deletion strains. GO enrichment, gene length, and type of promoter were all examined to understand if these genes were unique compared to all other genes. To perform the promoter-type analysis, we used the characterization of gene promoters as reported in (Basehoar et al., 2004). We also characterized several transcriptional phenotypes - antisense:sense ratio and pausing index around the TSS - for these frequently regulated genes in perturbation conditions. We examined these phenotypes split by the deletion strains in which the gene is differentially transcribed and those in which it is not, quantifying both at the gene level and split according to phenotype in the wild-type condition.

### Antisense transcription

For analysis of divergent antisense transcription at tandem genes, the sum of reads on the strand opposite the coding gene from 100 bp upstream of the sense TSS to 600 bp upstream of the sense TSS is divided by the sum of reads from the sense TSS to 500 bp downstream of the TSS on the coding strand. The genes selected were tandem gene pairs with each gene transcribed in the same direction and were obtained from (Churchman and Weissman, 2011). For analysis of antisense transcription, Pol II genes that did not overlap with another coding gene were chosen. The region analyzed spanned from the TSS to the poly(A) site as defined by taking the most abundant TSS and poly(A) site from (Pelechano et al., 2013). The sum of reads from the antisense strand was divided by the sum of reads from the sense strand. For all analyses, the log_2_ antisense:sense ratio was used. To generate antisense heatmaps, the log_2_ RPKM of NET-seq reads was used. Analysis at coding genes ranged from 250 bp upstream of the TSS to 4000 bp downstream of the coding TSS. To allow comparison between mutant and wild-type samples, a pseudocount of 1 was added to every position in all samples before calculating the log_2_ RPKM. Differential heatmaps were calculated by taking the log_2_ ratio of mutant / wild-type RPKM at each position. To assess the relative levels of antisense transcription across each gene in each strain, antisense and sense RPM were normalized by wild-type levels before the final normalized antisense:sense ratio was calculated. The “high-ratio” cluster of genes in the *set2Δ* and *rco1Δ* was identified by eye; gene length for genes in this cluster was compared to gene length of those genes not within this cluster using a Student’s t test. The relationship between antisense levels and gene length was quantified for each deletion strain using Pearson correlation (with Bonferroni correction for significance); this relationship was also visualized by binning genes according to length and quantifying mean antisense RPKM.

### Pausing index calculation

Pausing indices were calculated as the length-normalized Pol II density in the region of interest (−50 bp to +150 bp around TSS, ±100 bp around poly(A) sites, and ±10 bp around 5’ and 3’ splice sites) divided by the length-normalized Pol II density in the remainder of the gene, as illustrated in (**Figure 3A**).

### Metagene analysis

Only protein-coding, non-overlapping genes were included in the metagene analysis. The regions analyzed were −100 to +600 bp surrounding the most abundant transcription start site (TSS), −500 to +200 bp surrounding poly(A) sites, as identified in (Pelechano et al., 2013) and ± 25 bp surrounding annotated 3’ and 5’ splice sites. NET-seq signal across each region was normalized and the Loess smoothed mean (span = 0.01) and 95% confidence interval are plotted for NET-seq generated from each deletion strain across each region of interest.

### Splicing index calculation

Cac2Δ and wild-type RNA-seq data were retrieved from (Hewawasam et al., 2018) under the GEO accession number GSE98397. Splicing index calculations were determined for each gene by counting the number of reads that span exon junctions by at least 3 nucleotides and measuring the number of spliced reads divided by unspliced reads; splicing index = 2 * spliced reads / (5’ SS unspliced + 3’ SS unspliced reads) as in (Drexler et al., 2020).

### Extracting pause positions

Pauses were identified in previously annotated transcription units (Xu et al., 2009) of well-expressed genes (average of ≥ 2 reads per base-pair in two replicates). Pauses were defined as having reads higher than 3 standard deviations above the mean of the surrounding 200 nucleotides which do not contain pauses. Mean and standard deviation were calculated from a negative binomial distribution fit to the region of interest. Pauses were required to have at least 2 reads regardless of the gene’s sequencing coverage. Pauses were considered reproducible and used in downstream analyses when the Irreproducible Discovery Rate (IDR) is ≥ 1% between two replicates. To calculate the IDR of each pause, log_10_ of pause strength (number of reads in pause) for each replicate was used as a proxy for pause score. IDR was calculated using the est.IDR function of the idr R package (mu = 3, sigma = 1, rho = 0.9, p = 0.5) (Li et al., 2011). Reproducible pauses were visualized using the IGV genome browser (Robinson et al., 2011). Because the *cac1Δ*, *cac2Δ*, and *cac3*Δ strains were constructed by a different lab (Marquardt et al., 2014), these strains were excluded from these analyses. Additionally, *dhh1Δ* and *gcn5Δ* were excluded because of low sequencing coverage resulting in zero or 15 genes passing the coverage threshold, respectively.

### Pol II pausing location and strength

Pause density was calculated as the ratio of total number of pauses to the total length of the genome considered when extracting pause positions (combined length of all well-expressed genes in both replicates of each deletion strain). To identify deletions that induced similar pausing patterns, 8,816 pauses were found to be shared in at least eight strains and in regions sufficiently covered in multiple deletion strains. Shared pauses were visualized with a heatmap, clustered on both axes using the eisenCluster correlation clustering method in the hybridHclust R package (Chipman and Tibshirani, 2006), which takes into account missing data (where there was not enough coverage to confidently identify pausing in a particular deletion strain). Similarity in pause loci was also visualized as a scatter plot of the first two principal components. When calculating distribution of pauses across the gene body, all genes in which pauses were identified were normalized in length; the 5’ gene region was defined as the first 15% of each gene, the mid-gene region was defined as extending from the 15th percentile of gene length to the 85th percentile, and the 3’ gene region was defined as starting at 85% of gene length and extending to the annotated poly(A) site. The scrambled control for the pausing location analysis was created by randomly scrambling all identified pauses in all deletion strains across the gene in which they were discovered.

### Pol II pause loci sequence motifs

All analyses related to sequence motifs underlying pause loci were conducted using the MEME suite of tools (Bailey et al., 1994, 2009). The sequence ± 10 bp around each identified, reproducible pause (as well as the matched scrambled control) was extracted and used to run the MEME tool using parameters to find 0 - 1 motif per sequence, motifs 6 - 21 bp in length, and up to 10 motifs with an *E*-value significance threshold of 0.05 (Bailey et al., 1994). These significant motifs were compared to known transcription factor binding site motifs in the YEASTRACT_20130918 database (Teixeira et al., 2014) using the TOMTOM tool (Gupta et al., 2007) using default parameters, calling all hits as significant with an *E*-value greater than 0.1. TOMTOM searches were only performed on those motifs with a relative entropy greater than 5 and only the top match is reported.

### Random forest classifier for Pol II pausing loci

The predictive value of chromatin features for identifying Pol II pause loci was determined using a Random Forest model with the randomForest R package (Breiman, 2001). All reproducible Pol II pause loci were included in these analyses, as were an equal number of shuffled control loci. The shuffled control loci were selected to maintain the same number of real and control loci in each gene, controlling for effects of differential gene expression. In total, 51 chromatin features were compiled for all pause loci (**Table S5**) (Chiu et al., 2016; Oberbeckmann et al., 2019; Pelechano et al., 2013; Turner and Mathews, 2010; Umeyama and Ito, 2018; Vinayachandran et al., 2018; Weiner et al., 2015). Before applying the random forest classifier, we examined the distribution of values for each numeric feature (not discrete sequence) for real Pol II pauses compared to the scrambled control loci; statistical significance in the difference between these distributions was calculated with a Student’s t test, correcting for multiple hypothesis testing with the Bonferroni correction. From the random forest classifier, feature importance scores were generated using a random forest classifier with 75% training and 25% testing sets; for wild-type yeast, this is 10,495 training and 3,499 training loci. Due to the low number of reproducible pauses identified in the *dhh1Δ* and *gcn5Δ* deletion strains, they were excluded from these analyses.

Reported feature importance values are the mean decreases of accuracy over all out-of-bag cross validated predictions, when a given feature is permuted after training, but before prediction. Optimized parameters were selected for random forest classifiers trained using all features (**Figure S6B**):ncat = 4, mtry = 20, ntrees = 2500. ROC curve and AUC measurements were determined from binary prediction probabilities and calculated using the ROCR R package (Sing et al.). Prediction accuracy was determined by measuring the difference between the model’s predictions on a held-out test set and measured variables. The baseline score was determined using a “null” parameter that has the same value for every training and testing pair; thus, baseline represents the prediction accuracy with no additional information added to the model. To assess the transferability of a random forest classifier trained on Pol II pause loci in one strain, a model was trained on 100% of real and shuffled control Pol II loci from one deletion strain and then tested on all those pause loci in a second deletion strain, which was not included in the training set.

### Data and code availability

The accession number for the Illumina sequencing reported in this paper is Gene Expression Omnibus (GEO): GSE159603. All scripts and data analyses are available at https://github.com/churchmanlab/Yeast_NETseq_Screen. All plots were created in R using ggplot2 (Team, 2013; Wickham, 2016).

## SUPPLEMENTAL MATERIALS

Tables S1 to S6

### Supplemental Figures

**Figure S1.**
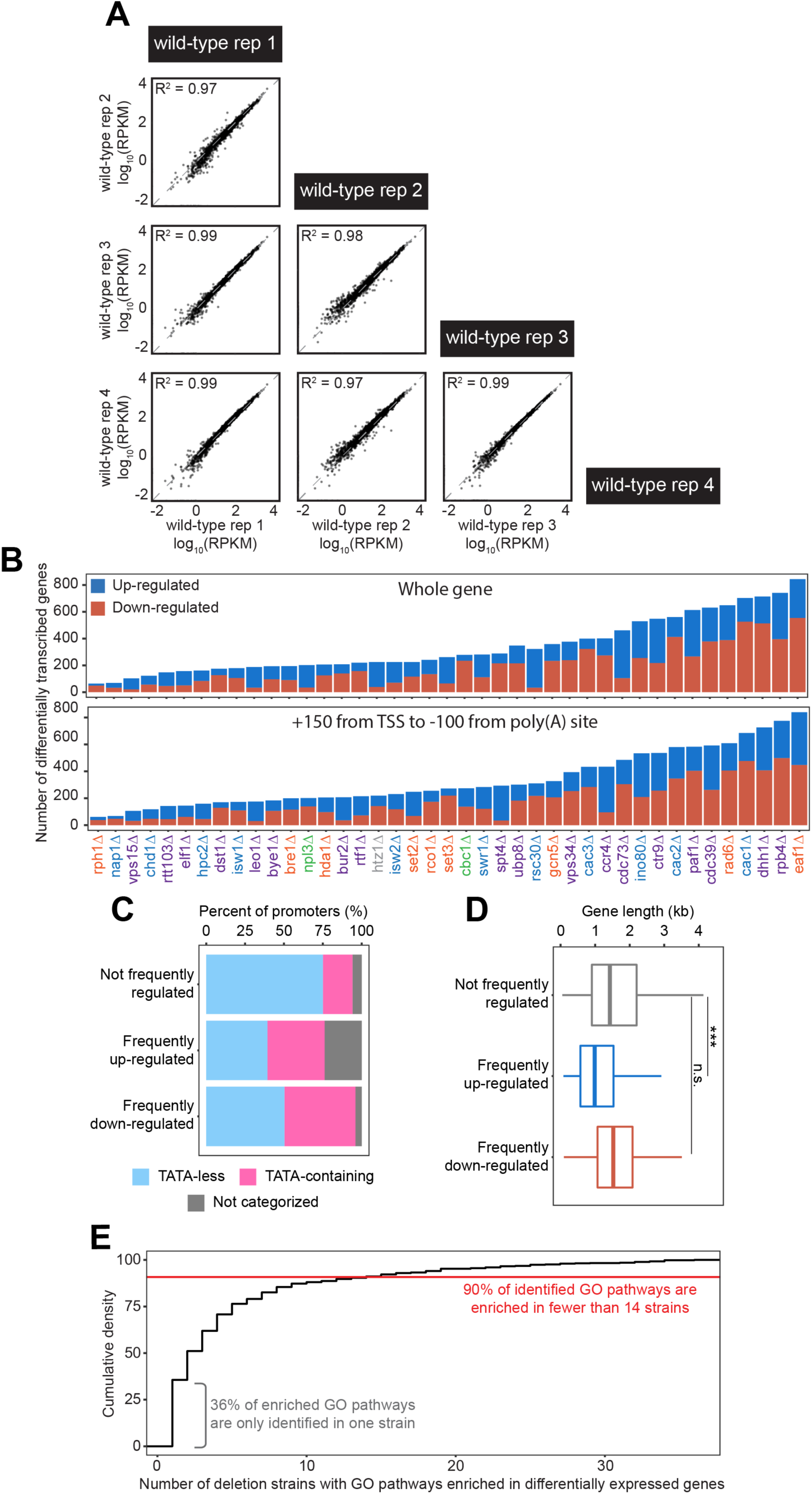
High-quality NET-seq screen data identifies largely different groups of genes with varying functions that differentially expressed across deletion strains. (A) Four biological replicates of the wild-type strain were used to establish baseline transcription activity. All replicates are highly correlated in gene RPKM by Pearson correlation. All four wild-type replicates are highly correlated. (B) Number of differentially expressed genes identified when using the entire gene and a sub-genic region to calculate expression. Regardless of whether the entire gene body (*top*) or a sub-genic region excluding pausing around the transcription start site and poly(A) site (*bottom*), there are similar numbers and trends across deletion strains in the amount of differentially expressed genes identified. Analyses conducted only on deletion strains with biological replicates. (C) Stacked bar chart illustrating the difference in proportion of genes that have TATA boxes in their promoters (*light blue*), are categorized as TATA-less (*pink*), or are not in either category (*grey*). Chi-squared test was used to quantify enrichment of TATA-containing promoters of frequently regulated genes. Gene promoters were categorized as in (Basehoar et al., 2004). (D) Distribution of gene lengths split by those genes that are not frequently regulated (differentially expressed in < 8 deletion strains, *grey*), frequently up-regulated (*blue*), and frequently down-regulated (*red*). Difference in distribution was assessed with a Student’s t-test. (E) The majority of genes found to be differentially expressed and the associated enriched GO pathways in one deletion strain are differentially expressed in multiple strains. Cumulative density plot illustrating that 40% of enriched GO pathways are only identified in one strain, with 90% of identified GO pathways enriched in 8 strains or fewer. There are several GO pathways that are enriched in 24 out of 31 deletion strains for which there were sufficient replicates to conduct differential expression analysis.

**Figure S2.**
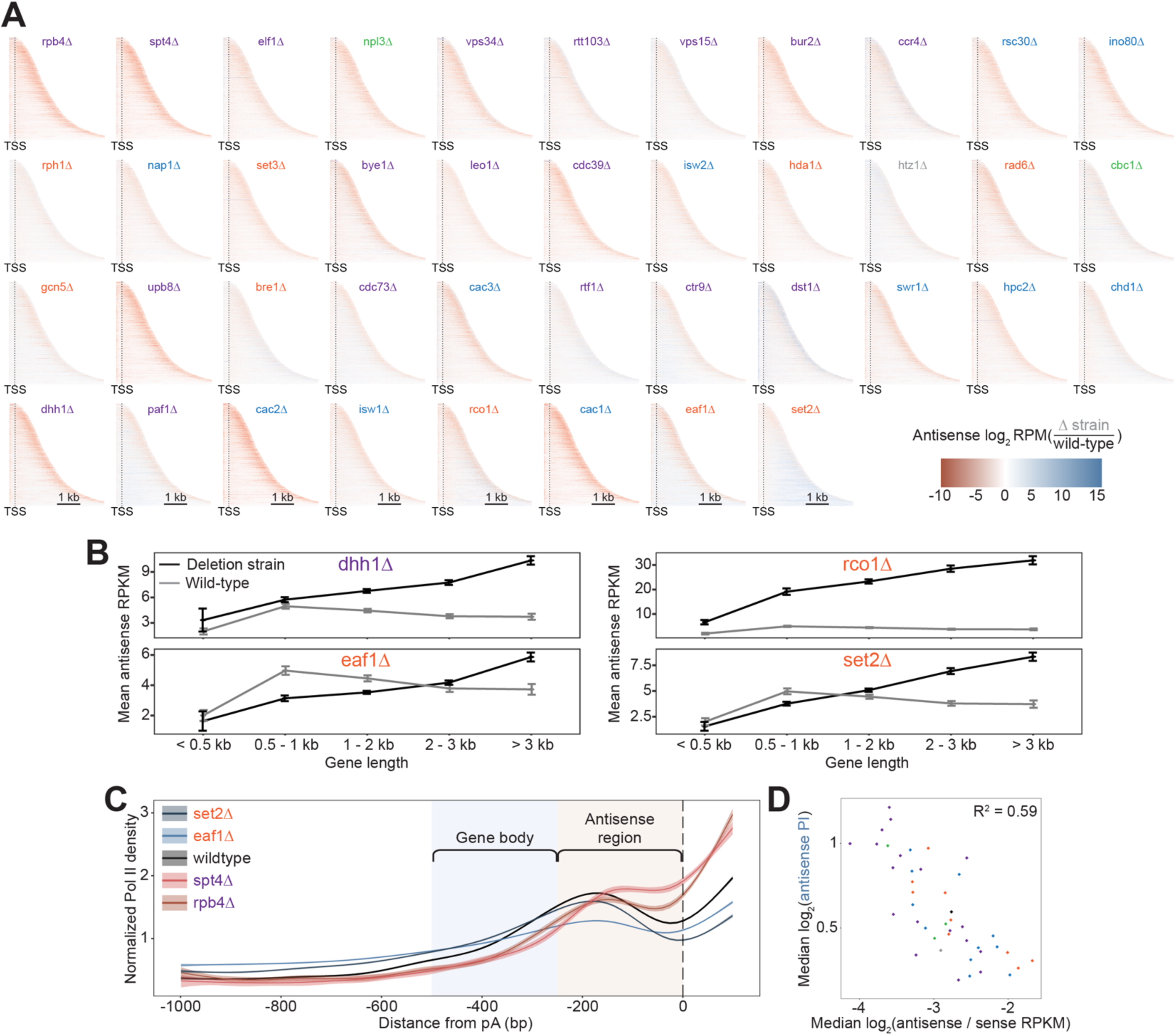
Antisense transcription is largely uncorrelated with gene length, and not uniformly distributed across gene bodies. (A) Heatmaps showing the relative enrichment of antisense transcription in deletion strains compared to wild-type across the gene body, from 250 bp upstream of the transcription start site to the end of the gene body, up to 4 kb downstream (ordered by gene length). All analyses conducted on non-overlapping, protein-coding genes (n = 3479). Heatmaps are ordered according to median antisense:sense transcription levels. (B) Line chart showing the gene-length dependence of antisense transcription levels, as measured by mean RPKM, in the *dhh1*Δ, *eaf1Δ*, *rco1Δ*, and *set2Δ* strains compared to wild-type. Error bars indicate standard error of the mean. (C) Metagene plot of antisense transcription from 1 kb upstream to the poly-adenylation sites of non-overlapping protein-coding genes for four deletion strains and wild-type. The gene body region is defined as 500 bp upstream of the poly(A) site to 250 bp upstream while the 5’ region of antisense transcripts is defined as 250 bp upstream of the poly(A) site to the poly(A) site itself. Deletion strains with high median antisense:sense ratios (*set2Δ*, *eaf1Δ*) have generally low antisense transcription in the defined 5’ region while deletion strains with low median antisense:sense ratios (*spt4Δ*, *rpb4Δ*) have high antisense transcription in this region. (D) Scatter plot showing the strong negative correlation between the median antisense:sense ratio and median 5’:gene body region antisense transcription ratios. Pearson correlation was calculated to demonstrate the strong negative correlation.

**Figure S3.**
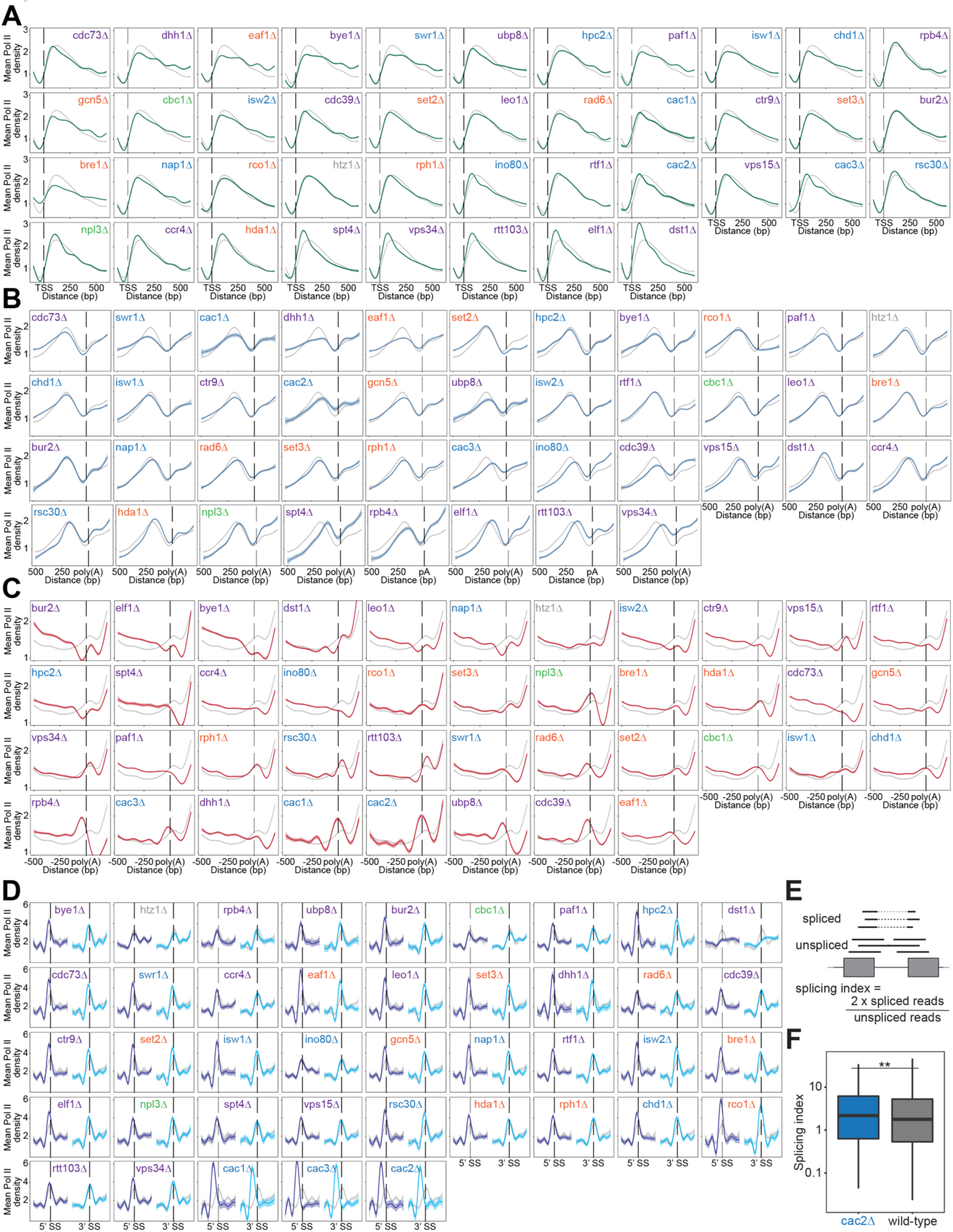
Heatmaps of Pol II density around RNA processing sites reveal differences in polymerase behavior across deletion strains, which can have functional consequences in specific deletion strains. (A) Normalized mean Pol II occupancy and the surrounding 95% confidence interval for −100 to +600 bp surrounding the most abundant annotated transcription start sites (Pelechano et al. 2013) (n = 2415 genes). Metagenes for each deletion strain (*green*) can be compared to the Pol II density in wild-type strains (*grey*). Deletion strains are ordered by median pausing index for the TSS region, as in Figure 3F. (B) Normalized mean Pol II occupancy and the surrounding 95% confidence interval for −200 to +500 bp surrounding the most abundant annotated poly(A) sites (Pelechano et al. 2013) (n = 2415 genes). Metagenes for each deletion strain (*blue*) can be compared to the Pol II density in wild-type strains (*grey*). Deletion strains are ordered by median pausing index for the antisense region, as in Figure 3G. (C) Normalized mean Pol II occupancy and the surrounding 95% confidence interval for the −500 to + 200 bp surrounding the most abundant annotated poly(A) sites (Pelechano et al. 2013) (n = 2415 genes). Metagenes for each deletion strain (*red*) can be compared to the Pol II density in wild-type strains (*grey*). Deletion strains are ordered by median pausing index for the poly(A) region, as in Figure 3H. (D) Normalized mean Pol II occupancy and the surrounding 95% confidence interval for the 50 bp surrounding annotated 5’ (*dark blue*) and 3’ (*light blue*) splice sites. Metagenes for each deletion strain can be compared to the Pol II density in wild-type strains (*grey*) (n = 252 genes). Deletion strains are ordered by median pausing index for the 5’ splice site region, as in Figure 3I. (E) Cartoon illustrating splicing index calculation. (F) Boxplot showing the distribution of splicing indices calculated in both the *cac2*Δ and wild-type strain. Significance was determined with a Student’s t-test. RNA-seq data was obtained from (Hewawasam et al. 2018).

**Figure S4.**
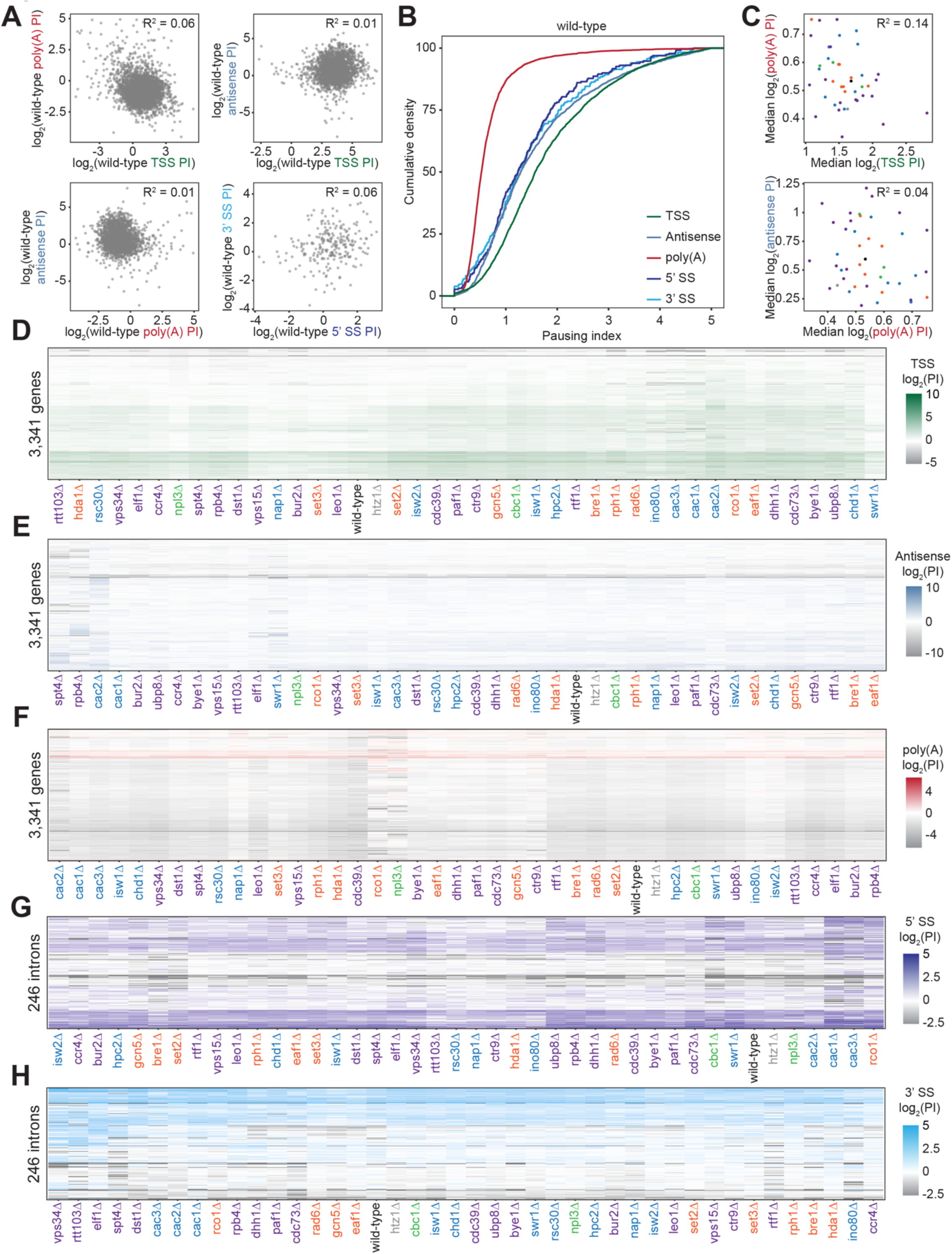
Pol II density is increased around RNA processing sites to varying degrees across deletion strains. (A) Scatterplot of the pausing index (PI) in the TSS and poly(A) region (*top left*), TSS and 3’ antisense (*top right*), poly(A) and 3’ antisense (*bottom left*), and the 5’ and 3’ splice sites surrounding introns (*bottom right*) for each gene in the wild-type strain. The lack of any relationship between these values is quantified by Pearson correlation. (B) Cumulative density plot illustrating the distribution of pausing indices for transcription start site (*green*), poly(A) site (*red*), 3’ antisense (*blue*), 5’ splice site (*dark blue*), and 3’ splice site (*light blue*) regions. In wild-type yeast, 25% of genes have a TSS PI ≥ 2.74; this PI value falls to 0.78 for poly(A) PI, 2.51 for 3’ antisense, and 2.18 and 2.35 for 5’ and 3’ splice site regions, respectively. Distributions of both splice site pausing indices are statistically the same, as determined by a Kolmogorov-Smirnov test (p = 0.273). (C) Scatter plot of the median pausing indices in the TSS and poly(A) regions (*top*) and poly(A) and 3’ antisense (*bottom*) for all deletion strains, colored as in Figure 1B. Relationship was quantified using Pearson correlation. (D) Pausing index for the transcription start site region across all non-overlapping protein-coding genes (n = 3,341). Both axes are hierarchically clustered, revealing genes with similar pausing densities as well as deletion strains that share pausing indices across their genomes. (E-H) Same as in (D), for pausing indices calculated across different gene regions - 3’ antisense, poly(A) sites, 5’ splice sites, and 3’ splice sites, respectively.

**Figure S5.**
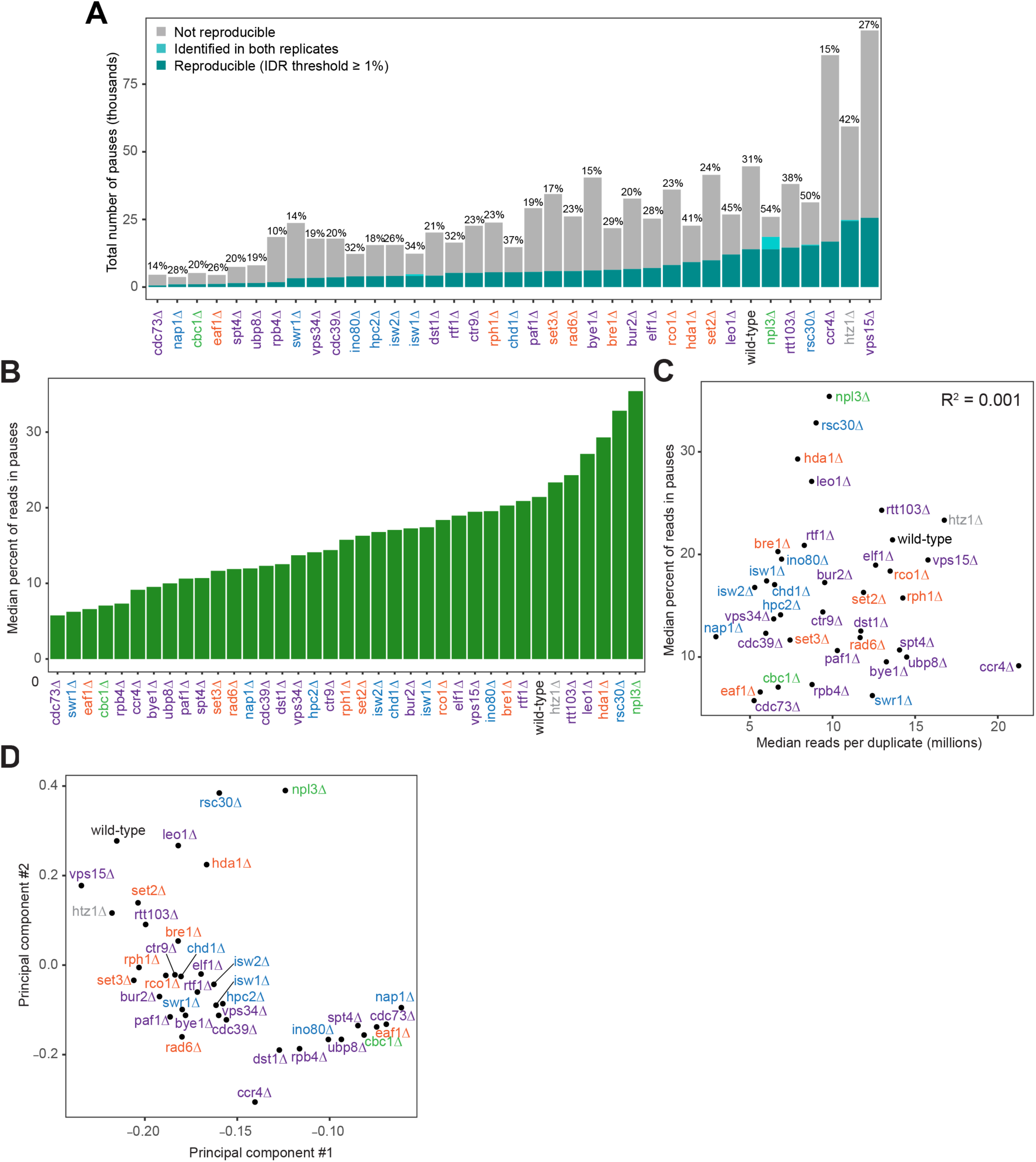
Pol II pausing behavior at single-nucleotide resolution across deletion strains reveal that pausing is balanced and dynamic in wild-type. **(A)** The number of reproducible pauses varies across deletion strains, as does the percent of pauses found to be reproducible. There is a median of 23% of pauses that reproduce across two replicates with an IDR threshold of ≥ 1%. Applying an IDR threshold of ≥ 1%, the strong pauses (*dark cyan*) are reproducible, while others do not meet this threshold (*cyan*), while still others are only present in one replicate (*grey*). Only genes meeting the coverage threshold for both replicates are considered by the pause-calling algorithm for each deletion strain. (B) Bar plot showing the median percent of reads, mapping to within highly-expressed gene bodies, contained within reproducible Pol II pauses, ordered from lowest to highest. (C) Scatter plot illustrating the relationship between the number of sequencing reads obtained in each duplicate for each deletion strain and the percent of NET-seq reads located in pauses across deletion strains. Relationship was quantified using Pearson correlation. (D) Principal component plot based on shared Pol II pause loci across the genome for different deletion strains. Deletion strains with more shared Pol II pause loci are closer together in this plot whereas deletion strains with very different Pol II pausing patterns are further apart.

**Figure S6.**
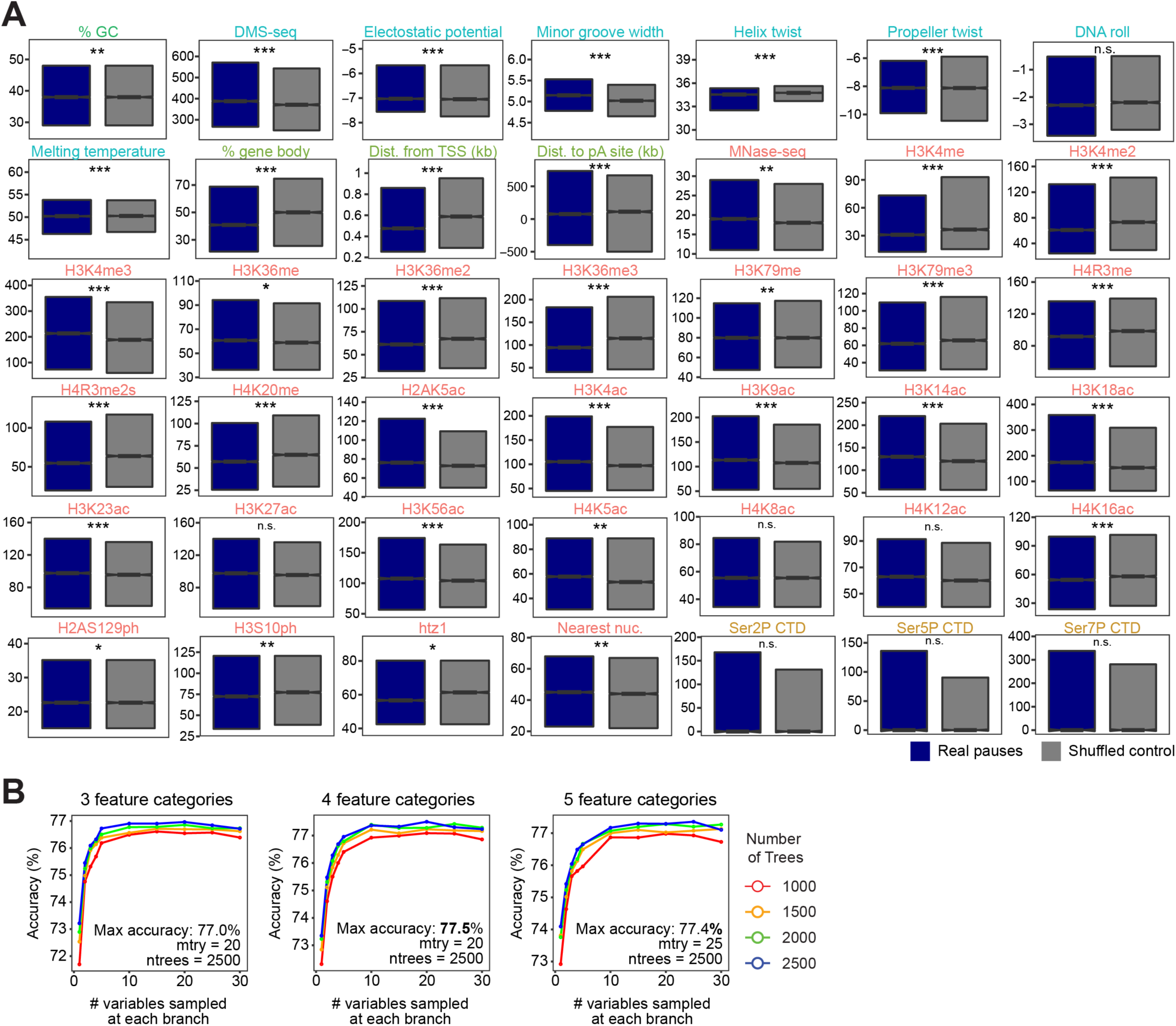
Chromatin features explain the location of some Pol II pauses in wild-type yeast using a random forest classifier. (A) Comparison in the distribution of values for each chromatin feature surrounding Pol II pauses in wild-type yeast (n = 13,994) compared to a shuffled control, in which the same number of pauses are shuffled, maintaining the same number of pauses within each well-expressed gene. Differences between the real and shuffled distributions were determined as significantly significant by a Student’s t-test with Bonferroni correction for multiple hypotheses. P-values are reported in Table S6. (* adjusted p-value ≤ 0.05; ** adjusted p-value ≤ 0.01; *** adjusted p-value ≤ 0.001). Colors correspond to legend in Figure 5E. (B) Accuracy of random forest classifiers trained to identify real and shuffled Pol II pause loci based on 51 features across parameter space. All continuous features were converted into categorical features by binning into 3 (*left*), 4 (*middle*), and 5 (*right*) categories of equal size. The number of variables randomly sampled at each branch (mtry) varied from 1 to 30 and the number of trees in the random forest classifier (ntrees) varied from 1000 to 2500. Parameters used for all downstream analyses were those that yielded the highest accuracy for each feature set (4 feature categories, 20 variable samples, and 2500 trees in forest for all features). All classifiers were trained on 75% of pause loci and tested with the remaining 25% of loci.

**Figure S7.**
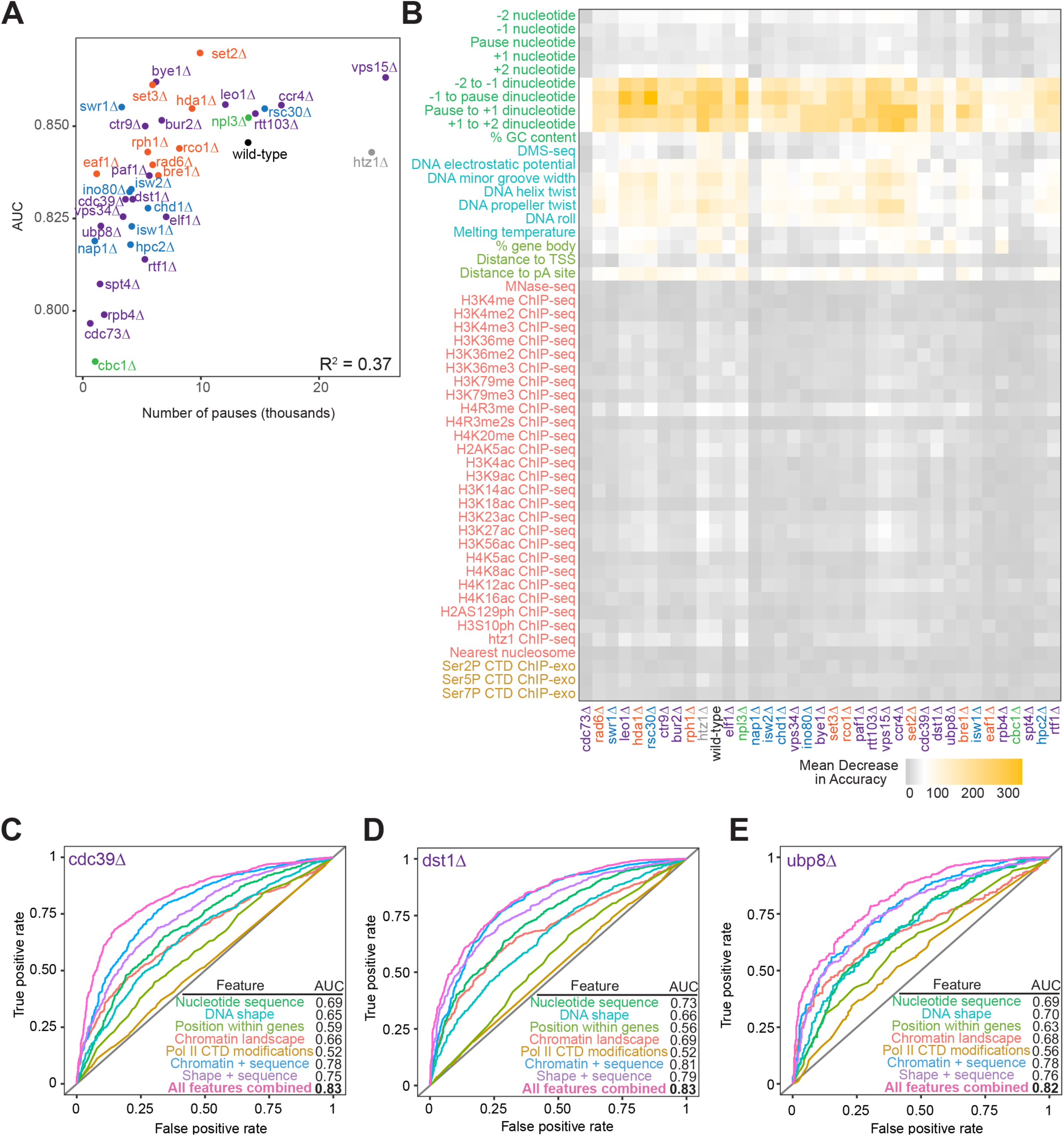
Random forest classifiers can predict Pol II pause loci across deletion strains, with different feature importance values across deletion strains. (A) Correlation between the number of reproducible pauses identified in each deletion strain and the AUC measurements for random forest classifiers trained on full set of chromatin features. The variation among deletion strain AUC measurements is not fully explained by the number of reproducible pauses identified in each deletion strain, as measured by Pearson correlation. (B) Heatmap illustrating feature importance for each feature, across all deletion strains. Deletion strains are hierarchically clustered along the x-axis, in the same order as in Figure 6A. (C-E) ROC curves and corresponding AUC values for random forest models trained on cdc39*Δ* (B) *dst1Δ* (C) and *ubp8Δ* (D), respectively.

## Notes

### Competing Interest Statement

The authors have declared no competing interest.

